# Optical mini-stroke of thalamic networks impairs sleep stability, topography and cognition

**DOI:** 10.1101/2021.08.25.457501

**Authors:** I Lenzi, M Borsa, C Czekus, T Rusterholz, C. L. Bassetti, C Gutierrez Herrera

## Abstract

Modelling stroke in animals remains a challenge for translational research, especially for the infraction of small *subcortical* arteries. Using combined fibre optics and photothrombosis technologies, we developed a novel model of optically-induced infarcts (Opto-STROKE). Combining our model with electrophysiological recordings in freely-behaving mice, we studied early and late consequent patho-physiological changes in the dynamics of sleep-wake circuits and cognitive performance. Here, focusing on inducing Opto-STROKE lesions in the intralaminar thalamus (IL), which in humans cause severe impairments of arousal, cognition, and affective symptoms, our model recapitulated important deficits on sleep disorders presented in humans including arousal instability, concurrent to an augmented slow-wave activity and a reduction gamma power bands during wakefulness. Moreover, during NREM sleep, spindle density was decreased and topographically shifted to frontal cortices when compared to control animals. Remarkably, gamma power and spindle density were correlated with decreased pain threshold and impaired prefrontal cortex-dependent working memory in Opto-STROKE mice relative to controls. Collectively, our combined method influences both anatomical and functional outcomes of the classical stroke procedures and offers new insights on the fundamental role of the media thalamus as a hub for the regulation of both sleep-wake architecture and cognition.

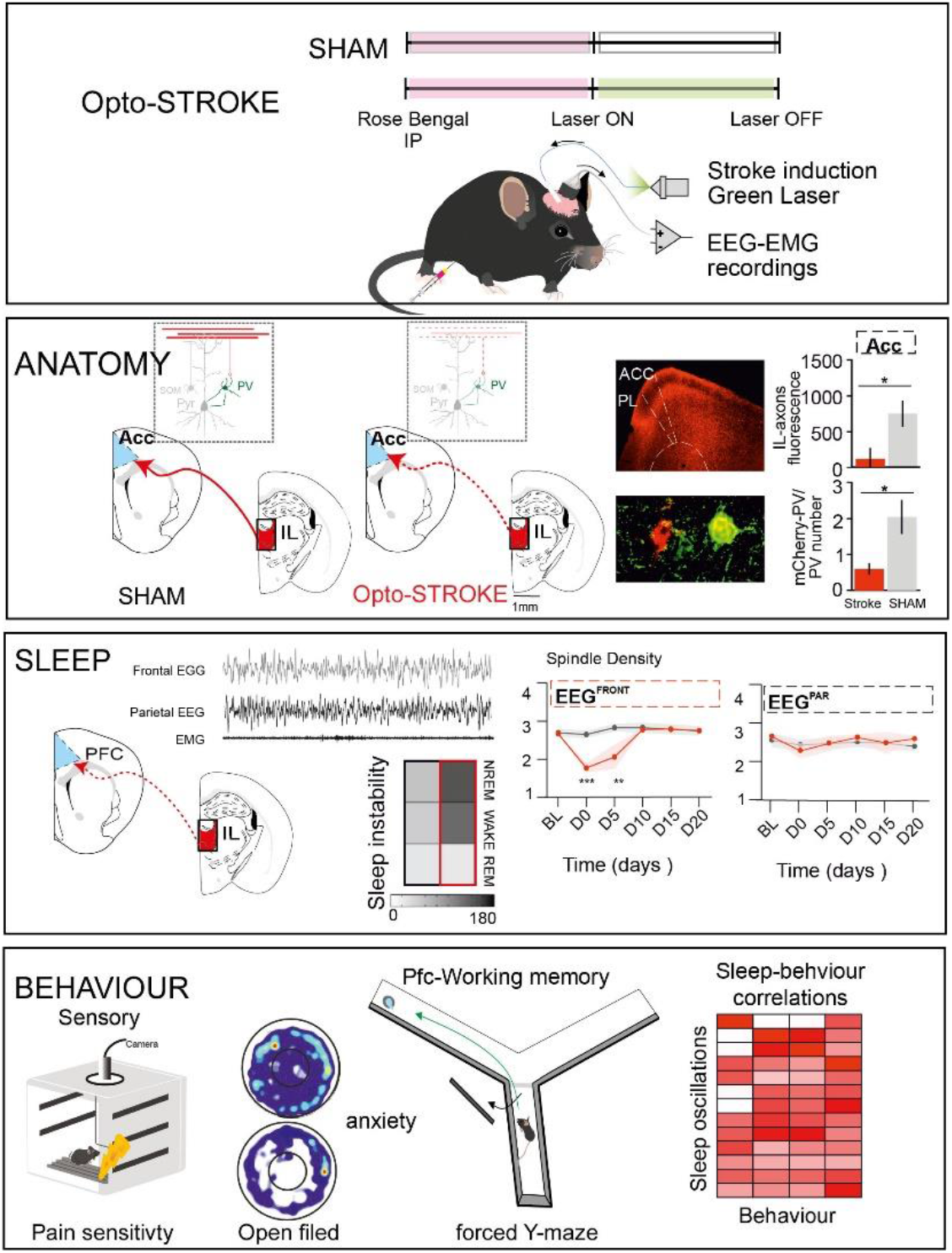

## Introduction

Stroke is a devastating brain disorder leading cause of morbidity and mortality worldwide that involves tissue damage and extensive structural modifications that are associated with severe physiological and behavioural consequences. The severity of clinical outcomes partly depends on the localization of the thrombus and the vascular territories involved. Although ischemic stroke management has advanced along with the management of pathological deficits, this disorder still remains a leading cause of long-term serious consequences in motor-, sensory-, sleep- and cognitive-related functions in patients which life quality never fully recovers. This is particularly true in thalamic infarcts involving the territory of the paramedian arteries (1). Paramedian strokes represent a unique category bearing a variety of outcomes, including altered conscious states, counting coma (2), hypersomnia with alterations in non-rapid eye movement (NREM) sleep architecture (3, 4); marked cognitive disturbances involving reference memory recall, attention, enhanced distractibility and confusion (5–7); dysfunctional emotional regulation, memory loss and perseveration (2); sensory processing disturbances, such as in pain sensitivity. While sleep-related dysfunctions often improve over time, however memory deficits and changes in behaviours tend to persist (2, 3). These numerous clinical symptoms suggest the insurgence of disconnection to subcortical structures. Despite major advances on acute stroke management and neuro rehabilitation, most of disabilities remain untreatable.

Paramedian strokes result from lesions of the tuberothalamic artery or, in rare cases, of the paramedian artery, irrigating the thalamic reticular nucleus (TRN), intralaminar thalamus (IL) which comprises the parafascicular nucleus (PF), medio-dorsal thalamus (MD) and central median thalamus (CMT), ventral anterior nucleus of the thalamus (VA), rostral part of the ventrolateral nucleus of the thalamus (VL), mamillothalamic tract, ventral amygdalofugal pathway, the ventral part of the internal medullary lamina, and the anterior medial (AM), ventral (AV) and dorsal (AD) thalamic nuclei (8, 9). Together as a functional class (2), these nuclei subserve arousal and various cognitive functions. Specifically, they are involved in the modulation of wake-sleep patterns, sleep oscillations, sensory processing, attention, goal-oriented behaviours and associative memory through their substantial connectivity to the cortex (thalamo-cortical), to other thalamic nuclei (intra-thalamic), as well as to other subcortical regions (10–16). According to their broad system of efferent and afferent projections, the IL have been recognized as higher order thalamic nuclei, involved in higher associative cognitive functions. Previous works have shown that MD and CMT neurons extensively project to parvalbumin expressing interneurons (PV+) in the anterior cingulate cortex (ACC) (17–20), which are responsible for feedforward inhibition ultimately modulating cortical excitability and brain states (21–23). Moreover, IL -prefrontal cortex (PFC) circuits have been shown to play a role in memory and attention-related tasks (24–26). Thus, understanding the relationship between the topography of sleep oscillatory activities and behavioural outcomes after stroke lesions is indispensable. Experimental stroke models have greatly advanced our understanding of development and consequences of stroke lesions. However, infractions of small subcortical arteries - one of the three major causes of human ischemic stroke (48) – still remains a challenge, as the absence of non-anesthetized freely-behaving models thus limiting the comparability of pre-clinical models to clinical cases of stroke (48, 49). These latter factors are particularly relevant in presence of vigilance disturbances, such as the ones caused by paramedian stroke. Here, we developed a novel model for optically induced mini-photothrombotic ischemia targeting subcortical areas (Opto-STROKE) that does not require anaesthesia and allows longitudinal (up to 4 weeks) multi-site electrophysiological recordings of sleep and circuit activity in freely behaving mice. By means of this novel technique, we studied the link between the topography of IL ischemic lesions and related changes in postsynaptic targets, sleep/wake and behaviour. Our findings showed that IL Opto-STROKE increases the fragmentation of sleep and wakefulness associated with sustained changes in slow wave activity and gamma oscillations in wakefulness when compared to SHAM control animals. Fronto-parietal expression of spindles was affected in IL-opto-STROKE animals in the acute phase after stroke. Interestingly, we found a negative correlation between sensory-related responses (pain) at day 20 after stroke and the power of gamma at the day of stroke induction, as well as between impairments in PFC-dependent memory retrieval in the subacute phase and spindle density on the acute phase post-stroke. In summary, our data implicates the IL as a fundamental hub for the regulation of both sleep-wake architecture and cognition, providing further insights on the relationship between sleep and cognition. Furthermore, this work contributes to the understanding of the temporal progression of thalamic strokes’ symptoms and plastic changes at the lesion site and within its post-synaptic targets. Overall, we propose the Opto-STROKE model as novel selective methodological approach to further dissect structural and functional consequences of local mini-strokes in subcortical structures and to explore crucial time windows for interventions.

## Methods

### Animals

C57BL/6JRj male mice (https://www.janvier-labs.com/en/fiche_produit/%20c57bl-6jrjmouse/10-20 weeks old, 23-30 g). Animals were kept in groups of 2-5 per individually ventilated cages (IVC) under controlled conditions (regular circadian cycle of 12:12 hours light: dark; lights on at 8:00 AM (ZT0), constant temperature 22-24°C and humidity 30%-50%). Animals included for sleep and behavioural experimentations were habituated to a light-dark cycle of 12 h (lights on at 08:00, ZT0) for 5 days prior the surgical procedures. Following viral transduction and chronic electroencephalogram (EEG) and electromyogram (EMG) electrodes implantation, mice were all housed individually and let recover for 7-10 days in custom-designed polycarbonate cages (300 mm x 170 mm). Then, animals were tethered and allowed to progressively adapt to the EEG/EMG and optic cables in their home cage for an additional 7-10 days to then remain plugged for the duration of the experiment. Animals were randomly assigned to two experimental groups: SHAM (control) and Opto-STROKE (experimental). Both groups underwent same injection, instrumentation and sleep and behavioural monitoring protocols. However, only stroke animals underwent the stroke induction protocol. All animals were treated according to Swiss animal care laws, and experimental procedures were approved by local authorities (Veterinary Office, Canton of Bern, Switzerland; license numbers BE 41/17 and BE 118/2020).

### Viral targeting

Viral injections were performed in animals of 10 to 16 weeks old as previously described (39, 50). Briefly, animals were anesthetized with isoflurane (4.0% induction, 1.5% maintenance). Body temperature was constantly monitored and kept at physiological range using a rectal thermo-probe and feedback-controlled heating system. Animals were fixed in a digital stereotaxic frame, analgesia was administered subcutaneously (meloxicam, 5 mg/kg), and lidocaine 2 mg/kg infused subcutaneously at the incision site. All animals received an intracranial injection of 200 nl AAV2-CaMKII-mCherry viral vectors (50 nl / min infusion rate) through 28 G stainless steel cannula (Bilaney), connected by a tubing to a 10 ml Hamilton syringe in an infusion pump (model 1200, Harvard Apparatus). Injections were performed to target the Intralaminar thalamus (IL, AP: −1.6 mm; ML: −0.75 mm; DV: −3.9 mm, 10°). Animals were given seven days to recover before instrumentation surgery.

### Instrumentation

Animals at 12-20 weeks of age were chronically implanted with a unilateral optic fibre (diameter 200 μm, part number = FT200UMT, ThorLabs) unilaterally above the IL (IL, AP: −1.6 mm; ML: −0.75 mm; DV: −3.8 mm, 10°) and bilaterally over the reticular thalamic nucleus (TRN, AP −0.8 mm, ML +1.7 mm, DV −3.5 mm) and the anterior dorsal thalamus (AD, AP −0.86 mm, ML +0.75 mm, DV −2.71 mm) together with an EEG / EMG connector (Straight Male PCB Header: cat. # 852-10-100-10-001101, Preci-Dip) as previously reported (39). Animals were instrumented with two screws over the frontal cortices (AP: 2 mm; ML: +/- 2 mm), two over the posterior cortices (AP: −3 mm; ML: +/-2 mm) and one the cerebellum as ground (Stainless steel screws diameter: 1,9 x 3,18 mm, Paul Korth GmbH & Co. KG) were in planted. In addition, two bare-ended EMG wires (Cat. # W3MUF 8/30-4046 55, 3 wire international (3WI)) were sutured to the neck muscles to record postural tone (Suppl. Figure 1A).

### Thalamic targeted strokes

Opto-STROKE induction was performed in animals at 13 to 23 weeks of age were intraperitoneally (IP) injected with 0.10 ml photosensitive dye Rose Bengal (Sigma Aldrich, cat No 632-69-9) diluted to 10 mg/mL in NaCl. Within 5-10 minutes after injection, 10 mW of 532-nm laser (LRS-0532-GFM-00100-03, Laserglow Technologies) was delivered through the optic fibre coupled to the laser into the AD, TRN or IL. For the SHAM procedure, animals received an IP injection of Rose Bengal but light was delivered. For anatomical investigation of at day 0, 5 and 10 of infarct induction, mice underwent the same procedure, but the optic fibre was removed from the brain 5 minutes post-illumination. All stroke inductions took place between ZT 4-5. Please note that the rage in the animals age was with the intent to model population variability and thus reduce discrepancies with the human disease.

### Data acquisition

#### Sleep recordings

EEG and EMG signals were amplified (× 1,000) using a multichannel differential amplifier (model 3500, AM System) and digitized at 512 Hz (NIDAQ 6363, National Instruments) using a sleep recording software (Sleep Score, View Point). A 48 hours’ baseline of spontaneous sleep-wake behaviour were recorded for all animals. Following, SHAM and Opto-STROKE animals were recorded in the infarct’s acute phase, or rather during stroke induction and for the following 12 hours (day 0), and semi-chronic phase at days five, ten, fifteen and twenty post-stroke, to follow the progression of sleep behaviours and oscillations.

#### Behavioural tests

Anxiety-, motor-, cognitive-related outcomes were assessed in sleep recordings-free days. Video recordings of the performances were analysed with Ethovision software (Noldus), allowing the tracking of the mice movements within the whole area and pre-determined fractions of the arenas, such centre in the open filed test. For the pain sensitivity testing sessions, videos were recorded via an USB-camera included in the fear conditioning system (Fear Conditioning 2.1, Ugo Basile).

### Sleep staging

As previously described (39, 50) electrophysiological data were manually scored and analysed using EEGlab (39). Exact transitions of three vigilance states were identified based on EEG/EMG frequency and amplitude. Wakefulness was determined by low-amplitude EEG and high amplitude EMG signals. Whereas NREM sleep was scored in period of high-amplitude EEG signals, rich in low-frequency oscillations (0.5-4 and 10-16) concomitant to reduced EMG tone. REM sleep was characterized by low amplitude, high EEG theta (5-9 Hz) power and isoelectric EMG with intermittent muscle twitches. Microarousals were scored and defined as cortical fast rhythm and EMG bursts of at least 1 s. Transitional states were defined as periods between Wake-NREM-Wake featured by the presence of low EMG power and EEGs including both slow and theta oscillations. Minimal transitional period length was defined as 5 sec.

#### Slow waves and spindles detection

Electrophysiological analysis was completed using custom written MATLAB scripts. Slow waves and spindles were detected via customised MATLAB scripts. For slow waves, adapted SWA-MATLAB detection toolbox (39, 53) was used to detect the negative envelope across the four EEG channels, filtered between 0.5 and 4 Hz, and consecutive zero-crossings were detected. The threshold in the amplitude of the detected event eliminates the potential individual differences on distances between electrodes and electrode depth that would affect the record amplitude. In a second pass, the activity over all four channels was examined for each SW detected on the negative envelope to obtain individual channel data. For spindle detection, a wavelet power was estimated between 10 and 16 Hz. Then wavelet functions classification was performed using the ratio between average power of spindle segments and spindle-free segments. The wavelet energy time series was smoothed using the 200 ms Hann window, and a threshold equal to 3 SD (SD: standard deviation) was applied above the mean to detect potential spindle events (54).

### Behavioural tests

All animals underwent three behavioural tests aimed to characterize the motor-, anxiety-, and cognitive-related differences between SHAM and Opto-STROKE animals see Supplemental Figure 1B-D. All tests were run at distinct times and days between sleep recordings. All measures were conducted at least 12 h apart from the last sleep recording in the light phase. To avoid a rewarding odour-based bias to mice choices, after each behavioural session, testing setups were carefully cleaned with 70% ethanol.

#### Open field test (OFT)

To quantify changes in exploratory tendency, motor integrity and anxiety, mice were placed one by one, in a round-shaped arena (60 cm in diameter) and let free to explore for 5 min (Supplemental Fig 1B). The OFT was repeated two times, once before stroke induction (OFT pre-stroke), and another time after (OFT post-stroke) (Suppl. Figure 1A). Measured outcomes were total time spent in the centre (s), total distance moved (cm) and mean velocity (cm/s) during the 5 min session.

#### Pain sensitivity test

Differences in sensitivity to foot shocks were assessed via a novel protocol using triplets of 2 s shocks of increasing intensity (0,05 mA - 0,6 mA) in 0.05 mA increments. Shocks with same intensity were interspersed with a 10 s interval (inter-triplet), while a 20 s time was used as intra-triplet interval. Pain responses to shocks were first defined, then categorized in 3 different pain thresholds (PT): absence of response, or “no response” (PT0); “backward movements” (PT1); “escape run”, “jump” and “immobility” (PT2) (modified from (51)). Scoring of the behaviours was done off-line.

#### Y-Maze alternation test (YM)

This test aimed to measure PFC-related cognitive performance (52). For habituation, mice were handled for one hour daily for one week prior to the beginning of the experiment. 3 days before starting of the behavioural test’s procedures animals were food restricted and monitored to maintain ~ 80% of basal body weight (Suppl. Figure 7). For the reward habituation, animals were provided with 0.1 ml of Sweetened condensed milk diluted 1:10 in water, that was later used as reward during training and testing in the YM. Animals were habituated to the Y maze arena (Figure 5H and Suppl. Figure 1C, top) on the second day of the food restriction period, where they were allowed to explore the maze for 10 min at ZT 3 and ZT 9. The food dispenser positioned in the three-arm ends were baited with food reward (0.1 ml of the diluted sweetened condensed milk). Then animals were trained for two sessions (T1 and T2) to alternate between left and right arms (“goal” arms) to find the reward (See Suppl. Figure 7D for learning curves). During this training sessions, goal arms were alternatively closed and baited to train the mice to run from a starting arm in the open goal arm and get the reward with a time limit of 1 min. Testing session consisted of 10 trials, during which animals were expected to remember the alternating strategy learned in the training sessions in order to get the reward. In the first sample trial (trial 0), one of the goal arms was closed, and mice were forced to get the reward in the opposite arm. In the next consecutive trials, mice were not prompt anymore with the door, and needed to alternate between right and left arm to find the reward. For a single trial, mice were given a maximum of five runs (or consecutive errors) interleaved by 30 s delay to get the reward. In the case that five consecutive errors occurred, animals were re-directed to the rewarding arm by closing the not-goal arm (see trial 0) to then proceed with a new trial. Performances were video recorded, and behaviour scored off line. Errors were ranked from E1 to E5 to evaluate choice perseveration (Suppl. Figure 1C, bottom). Finally, latency to the reward was measured to disclose possible motor deficiency in stroke animals.

### Immunohistochemical staining

Animals were sacrificed at the day of stroke induction (day 0) and post-stroke day 5, 10 and 21 with 150 mg/kg or 0,5 – 1 ml/kg pentobarbital intraperitoneal injection (Esconarkon ad us. vet., Streuli Pharma) and transcardially perfused with 1x phosphate-buffered saline (PBS) followed by 4% paraformaldehyde. Brains were postfixed for 24 hours, cryoprotected in 30% sucrose (48 h at 4°C), frozen in 2-methyl-butane on dry ice and cut into 40 μm sections. To measure the stroke volumes, every third slice was mounted onto a glass slide (the other two sets of sections were used for immunostaining), dried at room temperature, rehydrated, and processed for Nissl staining. Briefly, sections were immersed in Cresyl violet (Klüver Barrera, Bio-Optica), washed in distilled water and dehydrated in graded alcohols, cleared in xylene (Sigma Millipore), and mounted (Eukitt mounting medium, Bio-Optica) on gelatin-coated microscope slides.

The other two remaining free-floating brain sections were washed in 0.1% PBS with 0.1 % Triton A-10 (PBS-T) and incubated in blocking solution (1 h at room temperature; PBS-T with 10% donkey serum, Sigma Life Science). Then, sections were incubated with primary antibody to: mCherry (rat anti-mCherry, 1:1000, # M11217), microglia (ionized calcium binding adaptor molecule 1 (Iba1) (rabbit anti-Iba1 1:1000 Wako 019-19741), reactive astrocytes (mouse anti-glial fibrillary acidic protein GFAP, 1:800, Catalog # 13-0300), parvalbumin (rabbit anti-parvalbumin, 1:600, Ab11427 RRID:AB_298032) and NeuN (mouse anti-neuronal nuclei (1:500; cat. # ab104224; Abcam). Following repeated washes in PBS-T, sections were incubated with secondary antibodies (Anti-mouse Alexa Fluor 488; cat. # ab150113; Abcam; Goat anti-rat Alexa Fluor 555 (Cat. # ab150166); Goat anti-rabbit Alexa Fluor 488 (Cat. # ab150077); and Goat anti-mouse Alexa Fluor 647 (Cat. # ab150115) in PBS-T (2 h at room temperature). Finally, slices were washed in PBS 1X, mounted on microscope slides, and covered.

### Anatomical quantification

The extent of the Opto-STROKE lesion was evaluated by quantifying stroke edges delineated per section using ImageJ software (https://imagej.nih.gov/ij/). As described before (39), the lesioned area was measured in each brain slice and multiplied by the distance between sections to define the infarct volume.

Fluorescent signals from the immunohistological staining were performed by drawing regions of interest (ROI) normalized by the background fluorescence level (same size ROI) per individual section in ImageJ. A subset of brain sections was used to quantify the changes in neuronal projections to parvalbumin-expressing neurons in post-synaptic targets of the IL. For synaptic contact quantification, a squared area of about 250 um^2^ was used to delimit a fixed region in the ACC. Within the delimited zone, IL synaptic contacts (puncta) on to PV^+^ neurons were manually counted. After quantification, our measures were normalized by the number of PV+ neurons counted in the chosen area. Three to four sections per animal were analysed.

#### Microscopy

Images for anatomical analysis were acquired using a Nikon Eclipse Ti-E fluorescence microscope (M.I.C. facility - Mu40, CH-3008 Bern). For stroke volume quantification, images were acquired with a using an Olympus light microscope (Widmer Laboratory - Inselspital CH-3010 Bern) and magnified with 10x or 20x objectives. For the fluorescence immunohistochemical quantification, 10x and/or 20x magnification was used, and 60x for the quantification of IL-mCherry^+^ ACC-PV^+^ neurons.

### Statistical analysis

Differences in outcome parameters between SHAM and Opto-STROKE groups at each day of stroke progression were analysed using *two-way ANOVA* with multiple comparisons using LSM model (Prism 6 GraphPad; https://www.graphpad.com/scientific-software/prism/) and Post hoc Bonferroni testing for multiple comparisons between experimental groups and time points after stroke induction or otherwise as stated in the text and figure legends. All data are presented as mean ± SEM, and levels of statistical significance were set at threshold *P* < 0.05. For the analysis of the behaviour, paired and unpaired Student’s t test, or *two-way ANOVA* were used for the analysis between groups or within groups using Bonferroni post-hoc multiple comparison test. Animals that did not perform behavioural tests or lost the EEG/ EMG signals during longitudinal measurements were excluded from the analysis. Experiments were not conducted in a blinded fashion. Data were scored independently by two experimenters. At least 4 cohorts of animals were used for statistical analysis.

## Results

### Histological characterization of optical mini-stroke (Opto-STROKE) in thalamic regions

To overcome the limitations of current photothrombotic stroke models (49) and allow the study of subcortical strokes in awake behaving mice, we targeted different regions irrigated by the tuberothalamic artery to restrict the ischemic infarct to three discrete reticular and intralaminar (IL) functional thalamic class: the reticular thalamic nucleus (TRN), and the anterior dorsal thalamus (AD) and the IL (via the central median-thalamus (CMT)) (see methods). Quantification of the volume of ischemic infarcts across different stroke locations were consistent in size and restricted to the targeted nucleus and, to a lesser extent, to some of the adjacent nuclei (AD = 0.51 ± 0.06 mm^3^; TRN= 0.896 ± 0.109 mm^3^; IL 0.53 ± 0.23 mm^3^) (Figure 1A-F and Suppl. Figure 2 for TRN and AD and Suppl. Figure 3 for IL stroke). Labelling of GFAP and Iba-1 were used to quantify reactive astrocytes and microglia activation as markers of inflammatory pathological processes. We found a higher level of gliosis and microglia activation in Opto-STROKE animals as compared to SHAM at the lesion site (Suppl. Figure 2C and E). Quantification of the fractions of IL nuclei affected by the stroke revealed that the central median thalamic (CMT); dorsomedial thalamus (DMT); intermediodorsal (IMD); paracentral (PC); rhomboid (RH); and reuniens (RE) were affected in different proportions (CMT = 40.94 ± 15.70; DMT = 40.41 ± 8.97; IMD = 19.23 ± 7.37; PC = 19.83 ± 20.95; RH = 9.81 ± 4.68; RE = 13.26 ± 9.62) (Figure 1F).

**Figure 1.**
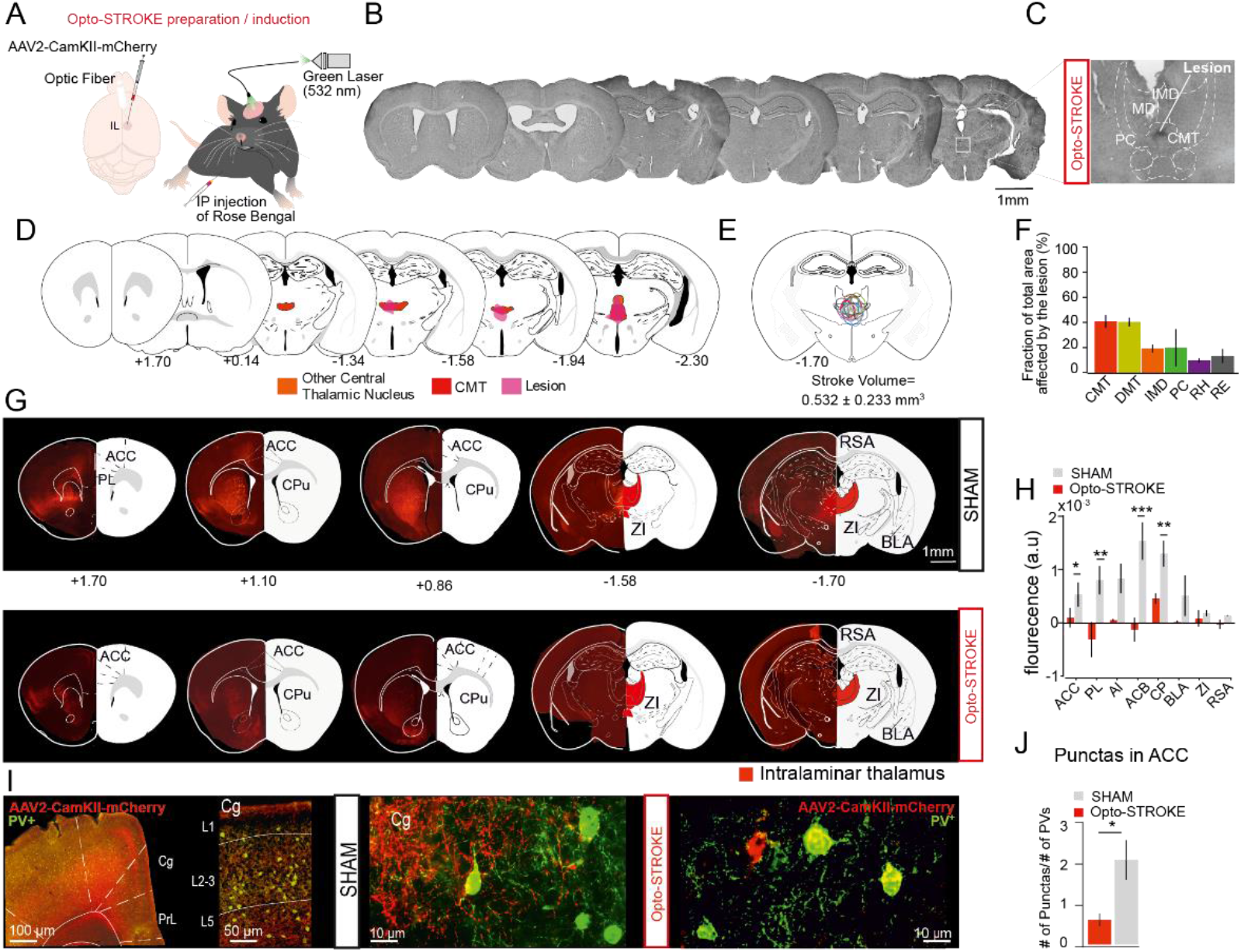
Characterization of Opto-STROKE lesions in the intralaminar thalamus (IL). (**A**) Schematic of animal model of targeted Opto-STROKE in the IL. (**B**) Representative AP distribution of IL lesion (Cresyl violet). (**C**) Close-up of IL thalamic nuclei showing the lesion site. (**D**) Schematic representation of anatomical atlas sections to (C) and coordinates from bregma (107). (**E**) Overlap of all obtained IL stroke lesions at coordinate −1.75 from bregma and mean lesion volume. (**F**) Bar graphs of the fraction (%) of IL thalamic nuclei affected by the lesion in Opto-STROKE animals (*n* = 9). Data are mean ± SEM. (**G**) Representative AP distribution of sections from SHAM (*top*) and Opto-STROKE (*bottom*) injected with AAV2-CamKII-mCherry. (**H**) Bar graph of the normalized mCherry fluorescence intensity in the target regions of IL in SHAM and Opto-STROKE animals (mCherry intensity in area/background fluorescence; SHAM (*n* = 3) vs Opto-STROKE (*n* = 4), (**I**) From left, in order: representative image of IL projections (mCherry, red) to prefrontal cortices (Anterior cingulate cortex (ACC) and prelimbic (PL)) and parvalbumin positive neurons (PV+, green); close-up of ACC cortex showing intense IL projections to ACC cortical layers; close-ups images from SHAM (*left*) and Opto-STROKE (*right*) animals showing IL-PV+ synaptic contacts. (**J**) Bar graph showing the quantification of the number of IL punctas on PV+ interneurons (Number of punctas/number of PV+ interneurons; SHAM (*n* = 4) vs Opto-STROKE (*n* =3), *two-way ANOVA* with Bonferroni post hoc test. Data are mean ± SEM. *Two-way ANOVA* with Bonferroni post hoc test; \**P* < 0.05, \*\**P* < 0.002, \*\*\**P* <0.0002).

Next, we characterized the structural changes of IL neuronal projections following Opto-STROKE using an anterograde tracing strategy (Figure 1G-H). To label IL neurons and their projections, AAV2-CamKII-mCherry viral vector was stereotactically injected in the IL area for stable expression of reporter fluorescent mCherry protein. Major IL neuron terminals were found in the anterior cingulate (ACC), prelimbic region (PL) of the frontal cortex, the anterior insular cortex (AI), zona incerta (ZI), caudo-putamen (CP), nucleus accumbens (ACB) and the basolateral amygdala (BLA) (Figure 1G). Interestingly, we found a significant decrease in mCherry fluorescence level in ACC, in Opto-STROKE compared to SHAM animals (ACC: SHAM = 843.89 ± 360.28; Opto-STROKE = −66.48 ± 419.70, unpaired *t-test*) (Figure 1H). To track IL circuit-specific synaptic changes, relevant for sleep modulation and cognition, we quantified IL synaptic contacts onto PV+ cell in the ACC after immunolabelling of GABAergic PV^+^ interneurons. Quantification of mCherry labelled post-synaptic contacts onto immunoreactive PV+ interneurons revealed a dramatic reduction in the number of IL puncta in Opto-STROKE as compared to SHAM animals (IL puncta: Opto-STROKE = 0.60 ± 0.31; SHAM = 2.05 ± 0.84; unpaired *t-test*) (Figure 1I and J). These results show: (1) specificity of the Opto-STROKE model in targeting restricted subcortical areas; (2) changes in the IL-target regions projection patterns following Opto-STROKE affecting particularly frontal cortices (PL and ACC); and (3) IL postsynaptic contacts on PV+ interneurons in the ACC were decreased post-Opto-STROKE.

### Intralaminar thalamic Opto-STROKE leads to rapid changes in arousal stability at the lesion onset

Previous clinical and fundamental investigations on the implication of thalamic lesions on regards to arousal, led to contrasting results with either loss (3, 4, 55) or no changes in arousability (56, 57). Here, we took advantage of the optical fibre technology for better temporal and spatial infarct resolution to investigate the functional consequences of IL Opto-STROKE. A remarkable feature of the Opto-STROKE model is the possibility to track changes from the onset (during) of stroke induction (5 min of laser on-Figure 3A) until later phases (here, up to 20 days) in un-anesthetised freely behaving animals. Animals were chronically implanted with EEG/EMG electrodes for characterization of their sleep-wake cycles (see methods and Figure 2A). Interestingly, we found that IL Opto-STROKE animals presented a decrease in the latency to the first stable NREM sleep (more than 50s long) immediately after stroke induction as compared to SHAM animals (SHAM = 1.07 ± 7.67 sec; Opto-STROKE = −13.55 ± 11.52 sec; unpaired *t-test*) (Figure 2B). Moreover, during the subsequent 24 hrs, sleep-wake macro-architecture was fragmented indicated by the increase of 56 +/ 14 % on the number of wake-NREM, NREM-wake transitions in OPTO-stroke mice relative to SHAMs (Figure 2C and Suppl. Figure 4A). Overall, analysis of the sleep architecture revealed a significant decrease in NREM sleep (SHAM = 0.57 ± 15.63 %; Opto-STROKE = −21.48 ± 17.66 %; unpaired *t-test*) (Figure 2D) and an increase in wake (SHAM = 6.94 ± 21.87 %; Opto-STROKE = 19.00 ± 19.00 %; unpaired *t-test*) as compared to SHAM animals on the subsequent 24 hours after Opto-STROKE.

**Figure 2.**
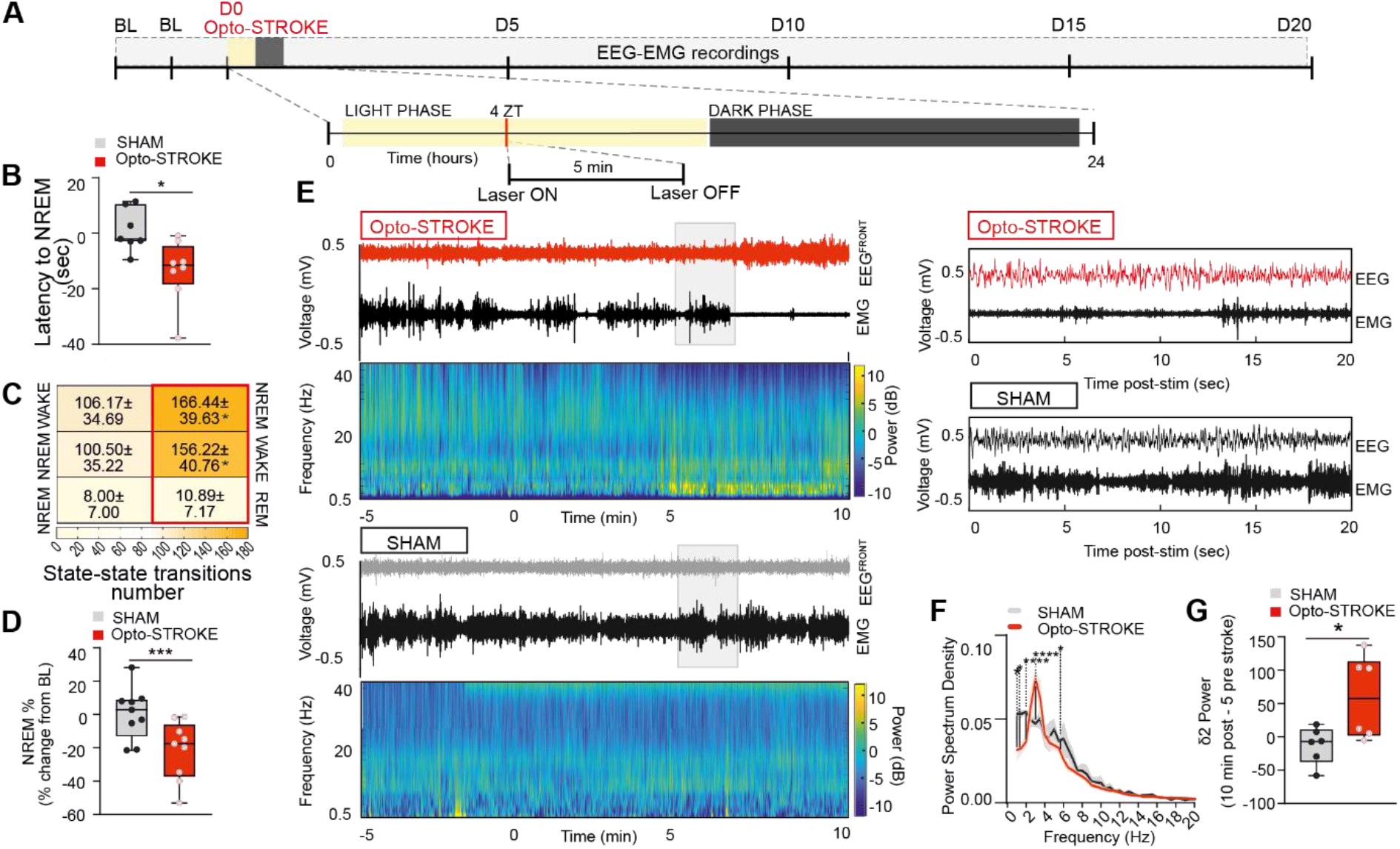
Changes in arousability immediately after IL Opto-STROKE induction. **A**) Experimental timeline for acute EEG-EMG recordings (D0). (**B**) Min-Max box-plots of the summary data of the NREM episode (min. 5 sec) onset latency (SHAM (*n* = 6) vs Opto-STROKE (*n* = 6), \**P* < 0.05, \*\**P* < 0.002, \*\*\**P* <0.0002), unpaired *t-test*. (**C**). Heat maps of WAKE-NREM-WAKE transitions’ level (Colour map light-dark yellow as increasing transition number, SHAM (*n* = 6) vs Opto-STROKE (*n* = 9), unpaired *t-test* \**P* < 0.05, \*\**P* < 0.002, \*\*\**P* <0.0002. \**P* < 0.05, \*\**P* < 0.002, \*\*\**P* <0.0002.) (**D**) Min–Max box-plots of NREM sleep % (SHAM (*n* = 9) vs Opto-STROKE (*n* = 9), unpaired *t-test*, \**P* < 0.05, \*\**P* < 0.002, \*\*\**P* <0.0002). (**E**) EEG-EMG traces and heat map of frequency analysis over 15 min (5 min pre-stroke induction; 5 min lase ON; 5 min post-stroke induction). (**F**) Power spectrum density of the frequency range 0-20 Hz within the time frame 5 min pre-stroke – to – 10 min post-stroke (SHAM (*n* = 6) vs Opto-STROKE (*n* = 6), unpaired *t-test, *P* < 0.05, \*\**P* < 0.002, \*\*\**P* <0.0002. (**G**) Min-max box-plots showing delta 2 (δ2) power between time 10 min post and 5 min pre-stroke (SHAM (*n* = 6) vs Opto-STROKE (*n* = 6), unpaired *t-test*. \**P* < 0.05, \*\**P* < 0.002, \*\*\**P* <0.0002). All data represents mean ± SEM.

To further characterized sleep/wake quality, oscillatory dynamics of cortical activity were investigated after Opto-STROKE induction. Time frequency analysis of the power 5 min before, during and after Opto-STROKE showed a shift towards lower frequencies with a significant augmentation in the power of the delta band (δ) (0.5-4.5Hz) following stroke in comparison to SHAM animals (SHAM = 0.10 ± 0.03; Opto-STROKE = 0.16 ± 0.03; unpaired *t-test*) (Figure 2E-F and Suppl. Figure 4B). This echoed a change in δ2 component (2.75-3.75 Hz) - a medio-dorsal thalamic indicator of sleep homeostasis (58) - restricted the frontal EEG cortices (EEG^FRONT^) (δ2^FRONT^ SHAM = −16.54 ± 33.38 %; Opto-STROKE = 18.96 ± 52 %; unpaired *t-test*) (Figure 2G), whereas no significant difference was found for the δ1 component (0.75-1.5) (Suppl. Figure 4B). The differences in δ1 and δ2 modulation after IL lesions were found to be persistent over the 20 days of sleep monitoring (Suppl. Figure 4).

### IL Opto-STROKE promotes arousal instability and reduces sleep efficiency

To characterize the semi-chronic effects of IL Opto-STROKE lesions on sleep-wake architecture and sleep oscillations, recordings were followed from day 0-20 from both experimental groups. Analysis was performed in ZT4-8 (light phase) and ZT16-20 (dark phase). Results showed that Opto-STROKE animals exhibited a general and marked acute fragmentation of NREM sleep and wakefulness in both the light and dark periods (number of NREM bouts % change from baseline light phase: SHAM = −6.17 ± 21.76; Opto-STROKE = 17.58 ± 16.65; NREM bouts dark: SHAM = 7.129 ± 13.395; Opto-STROKE = 42.471 ± 23.304; wake light phase (%) SHAM = −12.266 ± 18.695 %; Opto-STROKE = 8.858 ± 18.695 %; wake dark phase: SHAM = 1.96 ± 24.57; Opto-STROKE = 32.63 ± 24.95; unpaired *t-test*) (Figure 3A-D, and Suppl. Figure 5A), which tended to renormalize over time. Although, percentages of wake and NREM sleep were unchanged during the light period, lower amounts of wakefulness and increase in NREM sleep were present in Opto-STROKE mice during the dark period. REM sleep parameters were unaffected (Suppl. Figure 5 B and C).

**Figure 3.**
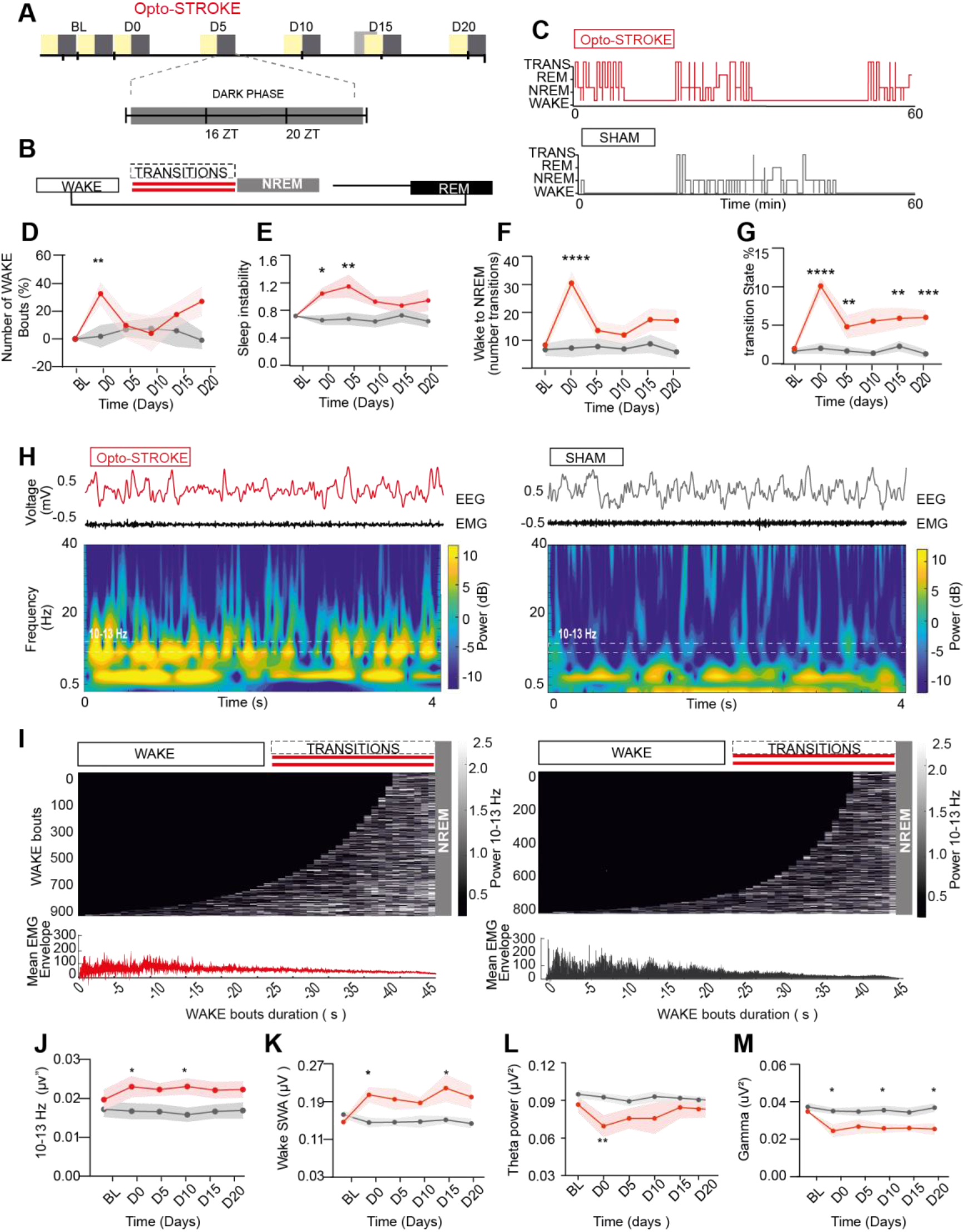
IL Opto-STROKE induces changes in sleep stability and efficiency. (**A**) Timeline showing sleep recording days and of 12 hours recording during light-dark cycle. (Hours analysed: light: 4-8 ZT; dark: 16-20 ZT). (**B**) Top to bottom: scheme showing natural transitioning between sleep states, with red connecting lines highlighting the WAKE-NREM-WAKE transitional periods enhanced in Opto-STROKE animals. (**C**) Hypnograms showing the increase in manually scored transitional states in Opto-STROKE animals (red) in comparison to SHAM (grey). (**D-G**) From left to right: wake number of bouts (SHAM (*n* = 9) vs Opto-STROKE (*n* = 10)) as change from baseline (in %), sleep instability (NREM bouts/WAKE bouts) (SHAM (*n* = 10) vs Opto-STROKE (*n* = 10)), transitional states number (SHAM (*n* = 8) vs Opto-STROKE (*n* = 11)) and % (SHAM (*n* = 8) vs Opto-STROKE (*n* = 11)) over time progression during the dark phase. (**H**) EEG-EMG traces and heatmaps of time-frequency analysis showing increased power in the frequency band between 10-13 Hz in Opto-STROKE animals during transitional states. (**I**) Stacked wake episodes at transition to NREM ordered from the shortest (5 sec) to longest (50 sec) in Opto-STROKE (left) and SHAM (right) animals, with respective mean EMG envelop (bottom), showing increase in 10-13 Hz at NREM transition and gradual decrease in EMG power (10-13 Hz power calculated normalizing each animal power within 10-13 Hz to the overall power in overlapping bins of 2 sec). (**J-M**) From left to right: line plots showing over-time progression of wake 10-13 Hz activity (SHAM (*n* = 9) vs Opto-STROKE (*n* = 8)), slow wave activity (SWA) (SHAM (*n* = 8) vs Opto-STROKE (*n* = 10)) and gamma power (SHAM (*n* = 8) vs Opto-STROKE (*n* = 10)) during the dark active phase. Statistical test. *Two-way ANOVA* with Bonferroni post hoc test was used as statistical test, \**P* < 0.05, \*\**P* < 0.002, \*\*\**P* <0.0002 were consider significant. Data is represented as mean ± SEM.

Further, a long-lasting increase on sleep instability was observed during the dark period following Opto-STROKE (NREM/wake bouts dark phase: SHAM = 0.67 ± 0.09; Opto-STROKE = 1.15 ± 0.043; *two-way ANOVA*) (Figure 3E). In contrast, during the light phase, Opto-STROKE animals showed small, but significant, increase in sleep instability at day 0 (NREM/wake bouts SHAM = 0.93 ± 0.075; Opto-STROKE = 1.18 ± 0.041; *two-way ANOVA*) (Suppl. Figure 3C). Unstable sleep was accompanied by a high number of wake-NREM-wake transitions (Number of transitions dark phase: SHAM = 7.25 ± 9.27; Opto-STROKE = 30.45 ± 13.60; light phase: SHAM = 12.63 ± 9.88; Opto-STROKE: 27.73 ± 14.34; unpaired *t-test*) (Figure 3F, Suppl. Fig 5D). Changes in wake and NREM sleep architecture progressively returned to baseline levels. However, sleep instability remained high over time.

The presence of wake and NREM sleep fragmentation during the animals’ active (dark) phase suggested potential further changes in arousal related activity. Indeed, spectral analysis of cortical EEG during wake episodes revealed a unique brain state signature featured by high amplitude 10-13 Hz oscillations that was accompanied by a decrease in the muscle tone, which was enhanced in Opto-STROKE relative to SHAM animals (Figure 3H-I). This state was further quantified using double-blind visual scoring to identify EEG with high 10-13Hz oscillation and low EMG power at wake-NREM sleep transitions which represented 6.84 ± 2.43 % of the total amount of vigilance states in Opto-STROKE as compared to 2.62 ± 0.29 % in SHAM animals. Remarkably, the quantity of these transitional state remained high across the light cycles for up to 20 days (Figure 3G, Suppl. Figure 5F and Suppl. Table 1). Moreover, additional analyses revealed an increased power in the 10-13 Hz frequency range in Opto-STROKE animals in comparison to SHAM during the duration of the experiment (Normalized power μV^2^ dark phase: SHAM = 0.02 ± 0.02; Opto-STROKE = 0.02 ± 0.001; *two-way ANOVA*) (Figure 3J and Suppl. Figure 6A).

Qualitative analysis of wake period indicated an enhancement of slow wave activity (SWA) across the light cycle (Normalized power (μV^2^) dark phase: SHAM = 0.15 ± 0.01; Opto-STROKE = 0.2 ± 0.024; light phase: SHAM = 0.15 ± 0.02; Opto-STROKE = 0.20 ± 0.03; *two-way ANOVA*) (Figure 3K). Remarkably, SWA changes were also restricted to the frontal EEG, as mentioned above for the acute changes (Suppl. table 2-3 and Suppl. Figure 6B). Furthermore, analysis of the wake spectral power during the dark-active phase showed a decrease in the theta (5-9 Hz) in IL Opto-STROKEs (Normalized power (μV^2^) SHAM= 0.09 ± 0.004; Opto-STROKE = 0.07 ± 0.01; *two-way ANOVA*) and gamma (30-60 Hz) power in comparison to SHAM control animals (Normalized power (μV^2^) SHAM = 0.04 ± 0.002; Opto-STROKE 0.03 ± 0.004; *two-way ANOVA*) (Figure 3L and M; Suppl. Figure 6C and D).

### Temporal and topographic renormalization of NREM sleep and oscillatory activities after IL Opto-STROKE

Paramedian strokes with inclusions of the IL have been reported to present a decrease in NREM sleep spindle density in acute phases, with a subset of the patients presenting long lasting spindle deficits (3, 4). Here, we sought to characterize the spindle oscillatory activities from IL Opto-STROKE animals in the acute and semi-acute phases following Opto-STROKE. First, we ran a spectral analysis of cortical EEG^FRONT^ and EEG^PAR^ signals during NREM for both light and dark phase. At day 0, we found a frontal-parietal dissociation of the sigma band (11-16 Hz) presented a decreased in EEG^FRONT^ power in IL Opto-STROKE animals (Normalized power SHAM D0 = 0.06 ± 0.01 μV^2^; Opto-STROKE D0 = 0.01 ± 0.01 μV^2^; *two-way ANOVA*).

Further, NREM sleep discrete spindle events were detected as previously reported (54). Remarkably, reduction in the spindle density was found in the frontal but not the parietal EEG derivation at day 0 and 5 (Spindle density (spindles/min): SHAM D0 = 2.65 ± 0.24; Opto-STROKE D0 = 1.77 ± 0.24; D5 SHAM = 2.83 ± 0.24; Opto-STROKE = 2.06 ± 0.62; *two-way ANOVA*) and to a minor extent during, the dark phase (Figure 4B).

**Figure 4.**
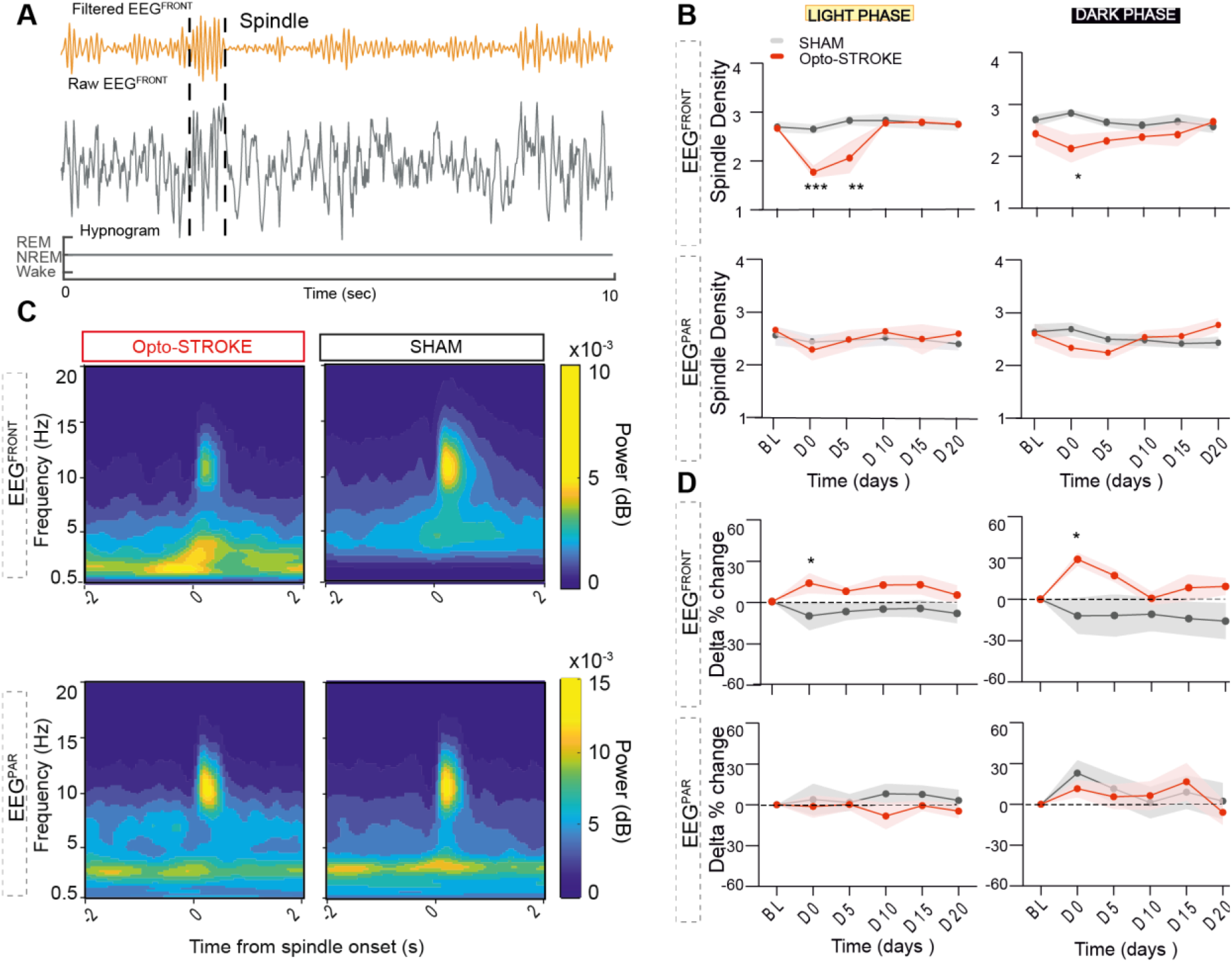
Spindle density and slow wave activity deficits renormalize after D10 post IL Opto-STORKE. (**A**) Representative traces of EEG signal filtered (top) raw (bottom) during NREM sleep for spindle detection. (**B**) Top: line plots showing EEG^FRONT^ sigma power and spindles density during dark phase (SHAM (*n* = 6) vs Opto-STROKE (*n* = 7), Bottom: line plots showing EEG^PAR^ sigma power and spindle density during dark phase (SHAM (*n* = 6) vs Opto-STROKE (*n* = 7). (**C**) Representative spectrograms showing coupling between spindles and slow wave activity in Opto-STROKE and SHAM animals (Left and right, respectively, upper panel). Bottom: Representative traces showing locking between delta and spindle activity. (**D**) Top: line plots showing increased EEG^FRONT^ delta power (% change from baseline) during both light and dark phase; bottom: line plots showing no change in EEG^PAR^ delta power in light and dark phase in Opto-STROKE animals (SHAM (*n* = 8) vs Opto-STROKE (*n* = 9). T*wo-way ANOVA* with Bonferroni post hoc test. Data are mean ± SEM. *P < 0.05, **P < 0.002, ***P <0.0002).

Expression of spindles has been linked to delta activity in humans, and rodents (54, 59, 60) as well as their temporal relationship with the expression of other oscillatory events (Figure 4A and C). Here, we found that in parallel to disrupted frontal spindle power, Opto-STROKE delta power was enhanced anteriorly (Normalized delta power all states light phase: SHAM = −6.01 ± 3.58 μV^2^; Opto-STROKE = 8.37 ± 5.27 μV^2^; dark phase: SHAM = −8.45 ± 4.51 μV^2^; Opto-STROKE = 18.36 ± 11.43 μV^2^; *two-way ANOVA*) (Figure 4C and D). Changes in delta power spontaneously recovered after day 5 post-stroke.

### IL Opto-STROKE increases pain sensitivity and impaired PFC-dependent working memory

The IL thalamus has been implicated in the regulation of stress and pain responses (2, 14, 61–64). To test this, mice underwent behavioural phenotyping for the assessment of locomotion/anxiety, pain threshold, and cognition-related responses (figure 5A). First, using an open field test (OFT), we observed a decrease in exploration (Figure 5B). However, we found no significant difference in the total distance travelled and the time spend in the centre of the open field arena either between SHAM and Opto-STROKE animals post -STROKE (Figure 5C-D), suggesting no impairments in the motor or stress related responses.

**Figure 5.**
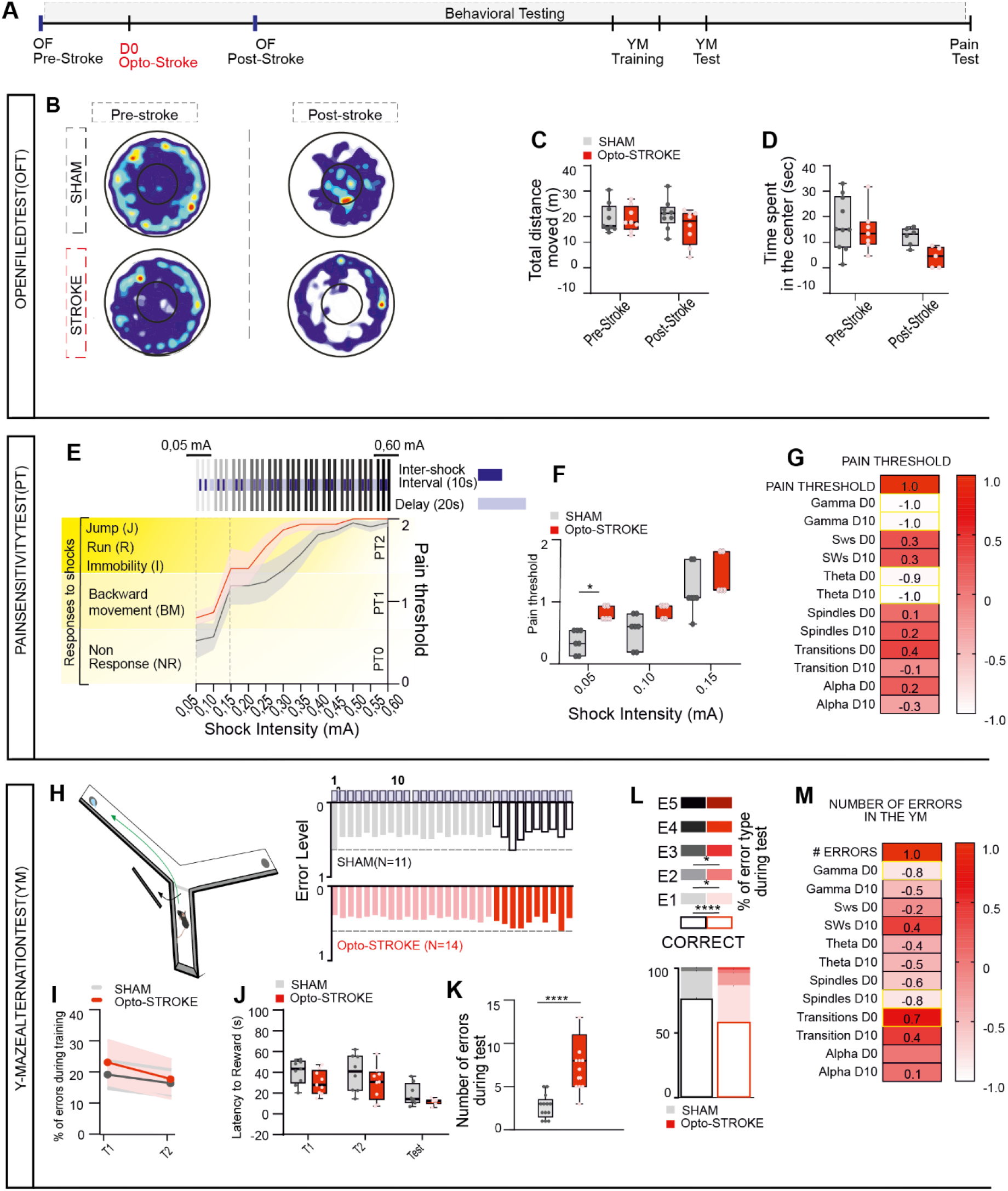
Stress-related responses, pain sensitivity and PFC-dependent working memory in IL Opto-STROKE. (**A**) Timeline of behavioural experimental procedures. Animals were tested before and post-Opto-STROKE induction (D0) in an open field (OFT) arena, and in the forced alteration task in the Y-maze (YM) and pain threshold test (PT) post-Opto-STROKE. (**B**) Representative heatmap showing motor activity of SHAM (upper row) and Opto-STROKE (lower row) mice at time pre- (*left*) and post- (*right*) Opto-STROKE. (**C-D**) Min-max box-plots showing total distance moved (C) and time spent in the centre (D) by SHAM (*n* = 10) and Opto-STROKE (*n* = 7) animals at time points pre- and post-Opto-STROKE induction. unpaired *t-test*. (**E**) Schematic representation (top) of the pain sensitivity test. Triplets of shocks (inter-shock interval: 10 secs; delay between triplets: 20 secs) with increasing intensity (0.05 mA-0.60 mA) were delivered to establish pain threshold in SHAM and stoke animals. Pain threshold to foot shocks was calculated via indexing behavioural responses (Non-response (NR): 0 (PT0); Backward Movements (BM): 1 (PT1); Jump (J), Escape Run (ER) and Immobility (3) (PT2)) and calculating average response within a triplet of shocks with same intensity. Bottom: line graph showing pain threshold in SHAM (*n* = 7) and Opto-STROKE (*n* = 5) miceand (I) Boxplot showing difference between groups in level of pain response to shocks of lower intensity (0.05 – 0.15 mA), unpaired *t-test*. (**G**) Heatmap of the correlation between pain threshold at 0.05 mA and sleep parameters (*Pearson correlation*, scale bar-1 < r < +1 (white= min value; red= max value). (**H**) Representation of YM set-up (left) and experiment structure with bar graph showing distribution of errors in SHAM (upper bar graph) and Opto-STROKE (lower bar graph) animals over sessions’ trials. Both training sessions (T1 and T2) and test session were composed by 10 trials interleaved by 30 s intervals. (**I**) Line graph showing percentage of errors accomplished during T1 and T2 by SHAM (*n* = 13) and Opto-STROKE (*n* = 15) mice. *Two-way ANOVA*. (**J**) Box-plots showing SHAM (*n* = 13) and Opto-STROKE (*n* = 15) animals’ latency to the reward during T1, T2 and test. unpaired *t-test*. (**K**) Box-plots showing number of errors made by SHAM (n = 13) and Opto-STROKE (n = 15) animals during testing session. unpaired *t-test*. (**L**) Stacked bar graphs showing level of error types in testing session (Error 1: E1; error 2: E1; error 3: E3; error 4: E4; error 5: E5) as a fraction of cumulative performance (total trials = 100%), with left legend indicating type of errors and significance. *Two-way ANOVA*, followed by Bonferroni post hoc test. (**M**) Heatmap of the correlation between errors number in the YM and sleep parameters (*Pearson correlation*, scale bar-1 < r < +1 (white= min value; red= max value). Data are mean ± SEM. \**P* < 0.05, \*\**P* < 0.002, \*\*\**P* <0.0002.

Then, to assess changes in pain perception, we used a pain sensitivity test (PT). A mild electric foot-shock test was employed with increasing shock intensity (0.05 mA-0.60 mA; Figure 5 E). Mice with IL Opto-STROKE presented lower pain threshold to shocks of lowest intensity as compared to SHAM animals (mean triplet pain threshold response level to 0.05 mA: SHAM = 0.38 ± 0.30; Opto-STROKE = 0.80 ± 0.18; unpaired *t-test*). Yet, no significant changes were found at higher shock intensities between groups (Fig 5F).

Lastly, to address the consequences of IL Opto-STROKE on learning and working memory, we used the forced alternation task in the Y-maze (YM). Mice were habituated, trained, and tested in the YM at day 12, 13 and 14 post-Opto-STROKE (Figure 5H, Suppl. Figure 1C, and Suppl. figure 7A). We found no significant differences in the number of errors during the training sessions or the latency-to-reward between the two groups (Figure 5I and J Suppl. Figure 7D), suggesting an absence of deficits in learning or motor performances. However, during testing, the total number of errors was significantly higher in Opto-STROKE as compared to SHAM animals (Total errors number: SHAM = 2.69 ± 2.84; Opto-STROKE = 7.73 ± 1; unpaired *t-test*) (Figure 5K). In fact, IL Opto-STROKE mice showed higher number of errors of type 1-3 with a significant lower percentage of correct trials compared to SHAM animals (Error level effect: *P* < 0.0001; interaction error level x group *P* < 0.0001, *two-way ANOVA*) (Figure 5L, Suppl. Figure 7C and Suppl. Table 4). Note that verification for potential reward-location preference was quantified. No preference was found in both experimental and control groups. (Suppl. Figure 7D).

To gain insights on the relationship between sleep and behaviour-related performances scores in OPTO-stoke mice, we performed a correlation analysis. Pain threshold was found negatively correlated with gamma power at both day 0 and 10 (Pain threshold-gamma D0: *r* = −0.983 *P* = 0.003; D10: *r* = −0.951 *P* = 0.049, *Pearson Correlation*) and theta power at D10 (Pain threshold-gamma D0: *r* =−0.935, *P* = 0.003; D10: *r* = −0.999, *P* = 0.005, *Pearson correlation*) (Figure 5G, Suppl. Figure 7E). Noteworthy, the number of errors in the YM was negatively correlated with gamma power (*r* = −0.809, *P* = 0.001, *Pearson correlation*) during the acute phase (D0). Furthermore, the long-term impairment in spindles had a negative correlation with working memory performance (*r* = −0.826, *P* = 0.006, *Pearson Correlation*) (Figure 5M, Suppl. Figure 7F). Collectively, these results are supported by previous studies showing the role of the IL in goal-oriented memory consolidation (21, 24, 62, 65–68). Importantly, our findings on the correlation between sleep and memory performance addresses the importance of alterations on sleep oscillations in the level of behavioural symptoms present at the onset and acute phases of stroke.

## Discussion

Our study introduced a novel model of Opto-STROKE in freely behaving mice, allowing the investigation of animals’ behaviour evolution from acute to semi-acute stroke phase. Here, we focused on subcortical IL Opto-STROKE lesions and related consequences on arousal and cognition. Our findings demonstrate the validity of such model via: (1) anatomical characterization, showing lesions to be limited to the intralaminar thalamus and stroke-related changes in inflammation and IL projection patterns (Figure 1 and Suppl. Figure 2 and 3); (2) acute sleep behaviour analysis revealing fast changes in arousability and stability (Figure 2 and 3); (3) over-time analysis of sleep architecture and oscillations showing stable fragmentation and increase in transitional states and progressive recovery (Figure 2 and 3); (4) study of cognitive impairments, revealing enhanced pain sensitivity and impaired working memory performance in a PFC-dependent task (Figure 5).

Current and widely-used stroke rodents models, including the middle cerebral artery occlusion (MCAO) and some photothrombotic ischemia, in large require a deep general anaesthesia and targets widespread areas, limiting the relevance of these translational approaches (44). Our photothrombotic Opto-STROKE model uses optical fibres (69) to precisely target sub-cortical brain areas and vascular territories in un-anesthetized freely behaving mice, allowing the monitoring of naturally occurring behaviours from stroke induction to acute and semi-chronic stages (up to 20 days). Importantly, plastic events within acute-stroke time window, and concurrent changes in sleep pattern and oscillations are key for intervention and recovery of cognitive, motor, and sensory functions (39, 44, 69), further supporting the relevance of our model.

Interestingly, we found that stroke lesions in the IL induced significant changes in the thalamo-cortical connectivity, and, in particular, in the IL-to- ACC and - PL circuits including synaptic contacts onto parvalbumin-expressing cells (Figure 1G, H, I, J). It is possible that some of IL projections contact other cell types in these areas, however, previous reports have shown that the majority of them project towards inhibitory interneurons (18, 26). Determining the exact contribution of IL-stroke to the changes in local cortical connectivity awaits further investigation.

Previous studies have indicated the role of the IL thalamic neurons in regulating arousal states and maintaining sleep stability (22, 61). Consistent with this, our results showed that IL Opto-STROKE is immediately followed by a decreased latency of the NREM sleep onset and increased sleep fragmentation, similar to clinical reports (3, 37, 70) (Figure 2B and C). This is accompanied by an overall increase in delta during the acute-stroke phase, with marked increase in the δ2 component in stroke animals (Figure 2G). These results are congruent with previous studies showing increased in δ2 following central median thalamic cell optogenetic inhibition (58), which mimicked post-sleep deprivation effects. Therefore, our results indicate a post-stroke sleep homeostatic need, which could be due to a change of sleep-dependent plasticity in the ischemic circuitry or circuit-related rearrangements, or both. Further confirmation of the former includes higher SWA during wakefulness, typically considered as a sign of increased sleep pressure (58). Yet this may also result from the high sleep and wake fragmentation in opto-stroke animals (Figure 3C and D). Overall, these results indicate a high level of sleep pressure and decreased ability to sustain arousal in IL stroke animals acutely, with a tendency to renormalize after 10 days post-stroke induction. This renormalization may harmonize contrasting results from other works on thalamic lesions and EEG recordings at semi-chronic stages, where no significant changes in the sleep or spectral power were observed (56, 57). Notably, our results provide evidence on the role of the IL in controlling the expression of slow oscillatory events in a topographic specific manner, which leads to their archetypal activity in the frontal cortex in healthy conditions, and to changes in such activity post-IL connectivity reorganization following Opto-STROKE.

A key feature of IL Opto-STROKE was an increased in transitional states, characterized by a 10-13 Hz frequency band (*alpha-like* activity) concomitant to a gradual reduction of the EMG power (Figure 3I)(Figure 3G and Suppl. Figure 5F). Interestingly, activity in the alpha-band has been hypothesized to reflect cortical activation, or more precisely, cortical excitation (71–73). For instance, during anesthetized states, the so-called “alpha-anteriorization” leads to a migration of alpha oscillations from the posterior cortex to the frontal cortex, particularly over the prefrontal cortex, in monkeys and humans (74–76).

At the cellular level, Lőrincz ML et al. (77) demonstrated that a subtype of excitatory thalamo-cortical neurons fire in burst at alpha frequency, driving inhibitory interneurons. Related to the IL Opto-STROKE, it may be that the reduction in IL excitatory connectivity to ACC PV+ cells (Figure 1J) is the underlying mechanism for the featured high 10-13 Hz power and the parallel increase in transitional states. Nonetheless, our results suggest a stroke-related reduction in cortical regulation and weakening of pyramidal neurons inhibition due to lack of IL input to PV+ interneurons (78–80), possibly leading to a higher cortical excitation. Notwithstanding early research (86) suggested a link between alpha-band activity to spindle activity and to thalamo-cortico-thalamic re-entrant loops (71), it is thought that alpha-band oscillations and spindle oscillations have a strikingly different physiological basis(81). Our results suggested that IL may be important for modulating the alpha band (72, 82).

Concomitant to changes in SWAs topography and alpha power, we found an acute decrease in sigma power and spindles density in IL Opto-STROKE animals (Figure 4). These results are consistent with human studies where reduction of spindles was observed after stroke (3, 4). Here, we reported evidence in favour of IL regulation of spindles and delta expression across the frontal cortex (54, 59, 83–85). Remarkably, we found a fronto-parietal dissociation of oscillatory activities where changes spindle density prominently affected the frontal cortices (Figure 4B-C) as reported for other thalamic nuclei such as the reticular thalamic nucleus (86–90). Future studies should concentrate on the understanding of how plastic changes occurring after stroke are related to changes in neuronal activity of distinct thalamo-cortical networks and their oscillatory activities from stroke progression’s acute to semi-chronic phases.

Previous studies in animals and humans have implicated the medial thalamus in memory and cognition (21, 25, 26, 33, 62, 67, 68, 91–93). Interestingly, in humans other brain disorders - characterized by strong deficits in cognition - have been often associated with altered PFC excitation including spindles density and changes in the structure and/or activity of the medial thalamus (19, 94–96). These investigations are further supported by the high density of medial thalamic projections to frontal cortical regions, while other thalamic nuclei preferentially regulate activity of more parietal cortices (18, 22, 25, 26, 61, 97). Here, we found a negative correlation between frontal spindles density and working memory performance that highlight the circuit specificity of our Opto-STROKE model and further supports the notion that these two phenomena might be functionally related and dependent on IL-PFC projections integrity (Figure 5M). Additional support comes from the found sensory-related deficits induced by IL Opto-STROKE, consisting in lower pain sensitivity threshold (Figure 5E-F), and impairments in recalling sequence of actions necessary to obtain reward and perseveration in a working memory task performed in the YM (Figure 5H-L). Our results further confirmed the importance of the IL thalamus in regulating both salient sensory stimuli processing and cognitive performance, possibly due to reduced central median-mediated PV+ inhibition onto pyramidal neurons in the PFC (17, 21, 23, 25, 26, 66, 91, 97–99).

Sleep- and arousal-related distinct oscillations, as slow waves, spindles, and gamma rhythms, has been shown to be beneficial in the recovery from traumatic brain injuries (39, 41, 69, 100–103). Moreover, previous studies have shown that sleep-related neural rhythms regulate synaptic plasticity and memory consolidation, while their enhancement is beneficial for both neurological and psychiatric improvements (41, 104–106). In this context, our un-anaesthetised mini-stroke model will be of particular interest since it enables the anatomical and functional dissection of specific brain areas in variable and complex pathological condition such as stroke. Thus, this study provides important insights about sleep- and arousal-dependent circuit activities and its relation to sensory and cognitive processes. Ultimately, it may open new ways of future circuit- and/or region-specific therapeutic strategies and improved personalized treatment.

## AUTHOR CONTRIBUTIONS

Author contributions: C.G.H. and I.L. conception and design of research; I.L and M.B. performed experiments; T.R. wrote and adapted Matlab custom scripts for data analysis, I.L., M.B. and C.K. analysed data; C.G.H. and I.L. interpreted results of experiments; I.L. and C.G.H. prepared figures; I.L and C.G.H drafted manuscript; C.G.H, I.L., M.B. and C.K edited and revised manuscript; C.G.H. approved final version of manuscript.

## Acknowledgements

We thank the Zentrum for Experimental Neurology for hosting our research and providing with technical support. To Prof Widmer for the use of the fluorescence microscope and the imaging facility of the University of Bern (Mu40). This work and I.L. were supported by University of Bern Interfaculty Research Cooperation “Decoding Sleep” (Gutierrez Herrera, C.); C.K and T.R were supported by the Inselspital University Hospital (Gutierrez Herrera, C), the University of Bern. M.B. work was part of the Biomedical Sciences Master program at the University of Fribourg-University of Bern.

