## Supplementary material for "Optical mini-stroke of thalamic networks impairs sleep stability, topography and cognition": Supplenetal figures

**Supplemental Figures**

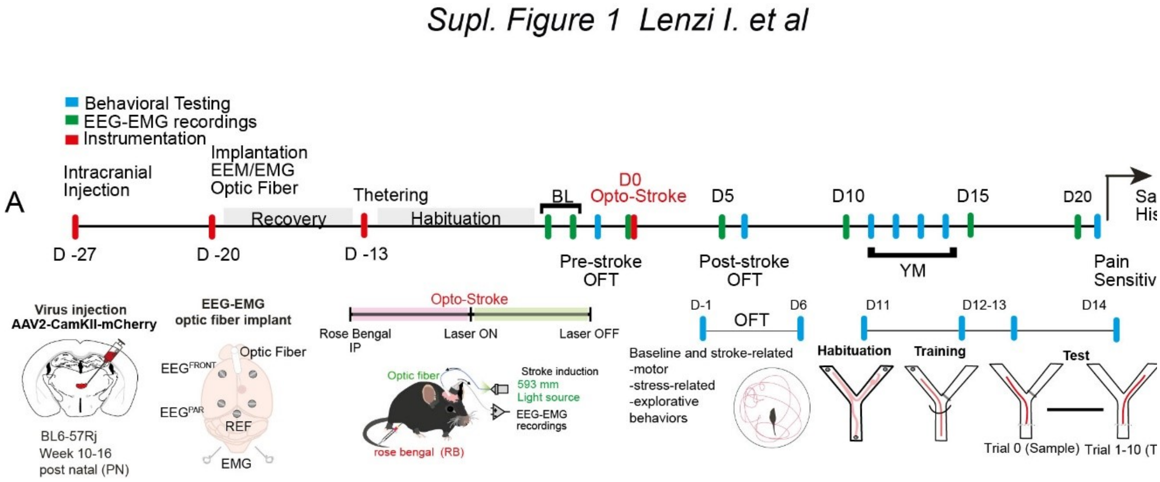


**Supplemental Figure 1. Timeline of experimental sleep and behavioural testing procedures.** (**A**)**.** Timeline showing temporal order of experimental procedures. Briefly, C57BL/6J animals underwent surgical preparation including intracranial injection of AAV2-CamKII into the IL and after one-week implantation of an optic fibre in the IL and 5 screws: 2 frontals and 2 parietal EEGs (respectively $EEG^{FRONT}$ and $EEG^{PAR}$), and 1 as reference over the cerebellum. 2 additional wires were implanted at the neck muscle (EMGs). After, mice were left recovering for 7 days and then tethered for habituation for 10 days. Later, sleep baseline (BL) recordings were acquired for 48 hours. Before stroke induction, mice underwent testing in the open field (OFT). The following day, stroke was induced in the IL via IP injection of rose Bengal and green light (532 nm) delivered through the optic fibre, while EEG/EMG were recorded simultaneously. Sleep recordings were acquired at day 0, 5, 10 15 and 20 post-strokes. Post-stroke testing in the OFT was achieved at day 6 post-stroke. YM habituation, training and test were run between day 11-14 post-stroke induction. At day 21 post-stroke mice underwent pain sensitivity test, before being sacrificed. Brains were collected for histological characterization of stroke.

*Suppl. Figure 2. Lenzi I. et al*

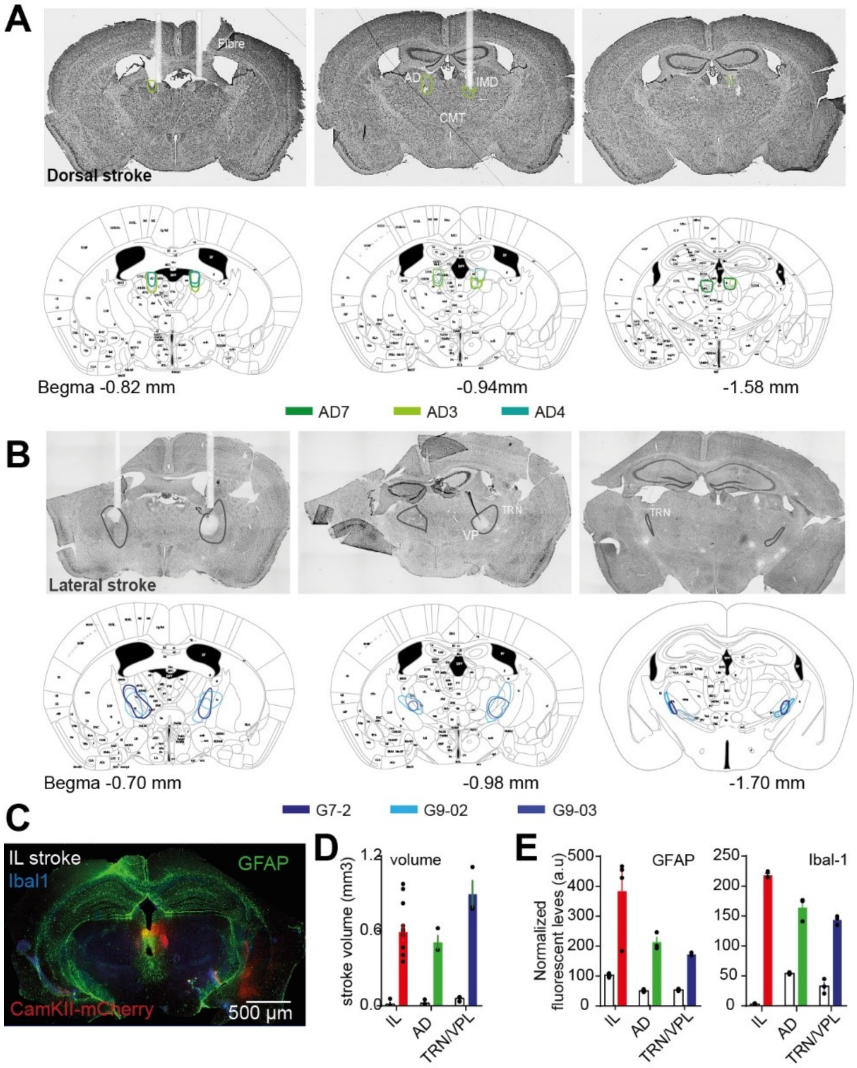


**Supplemental figure 2. Validation of targeted thalamic Opto-STROKE model and anatomical characterization.** (**A**) Representative images of Cresyl violet (CV) stained brain sections showing Opto-SROKE lesions in the anterior-dorsal (AD) thalamus (top) and correspondent atlas showing summary of lesions per animal (n=3). (**B**) Opto-STROKE lesion in the and reticular thalamic nucleus (TRN) and ventral posterolateral thalamus (VPL) on the top. Correspondent atlas showing the lesion distribution between animals (bottom). (**C**) Representative image showing level of inflammatory markers in an animal with Opto-STROKE lesion in the CMT (Green: Glial fibrillary acidic protein (GFAP); Blue: ionized calcium-binding adapter molecule 1 (Iba-1). (**D**) Bar graphs showing volume quantification of stroke lesions in the IL ($n$ = 9), IL ($n$ = 3) and TRN-VPL ($n$ = 3) (Data are mean ± SEM, unpaired *t-test*); (**E**) From left to right: bar graphs showing level of GFAP fluorescence in the IL, AD and VD-TRN; bar graphs showing level of Iba-1 fluorescence in the IL, MD and TRN-VPL (SHAM ($n$ = 3) vs CMT ($n$ = 4) vs AD ($n$ = 3) vs TRN-VPL ($n$ = 3) Opto-STROKE, unpaired *t-test.* Data are mean ± SEM. *$P$ < 0.05, **$P$ < 0.002, ***$P$ <0.0002).

*Supplementary Figure 3. Lenzi I. et al*

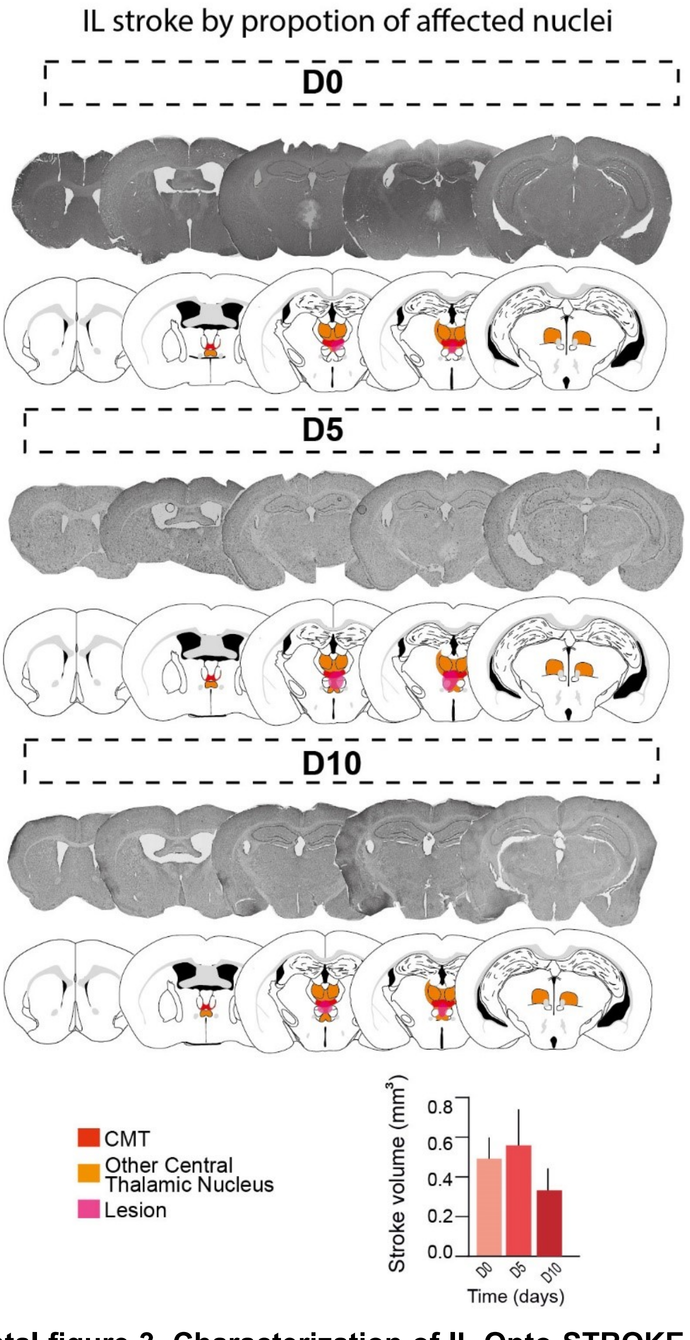


**Supplemental figure 3. Characterization of IL Opto-STROKE lesion over time. (A)** Anterior-posterior distribution of Opto-STROKE lesions in the IL at D0, D5, and D10, up, central and bottom panels, respectively. On the bottom, the correspondent atlas sections with coordinates and schematic of lesion proportions of affected nuclei (Red: Centromedian thalamus (CMT); Orange: other intralaminar nuclei; pink: lesion). **(B)** Bar graph showing volume of the lesions at D0, D5 and D10 (D0 vs D5 vs D10, (*n* = 2), bar graphs. Data are mean ± SEM).

*Supplementary Figure 4.Lenzi I. et al*

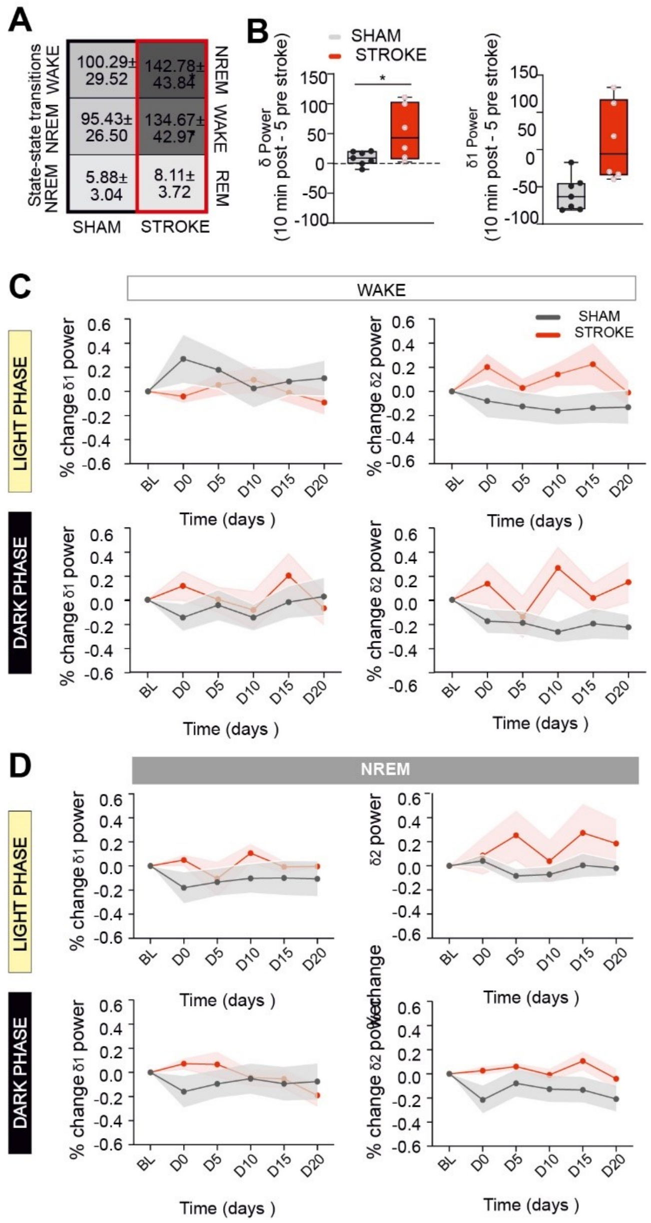


**Supplemental figure 4. Changes in arousability following Opto-STROKE in the IL are characterized by increased transitions and delta power in wake.** (**A**) Heatmap describing increase in WAKE-NREM-WAKE transitions in Opto-STROKE animals (Colour map: light-to-dark grey as increasing transition number, SHAM (n= 8) vs Opto-STROKE (n=10), unpaired *t-test*. Data are mean ± SEM, $*P < 0.05$, $**P < 0.002$, $***P$ <0.0002). (**B**) Min-max box-plots showing changes in delta (δ) and delta 1 (δ1) power from 5 min pre to 10 min post-stroke induction (SHAM ($n$ = 6) vs Opto-STROKE ($n$ = 6), unpaired *t-test*, $*P < 0.05$, $**P < 0.002$, $***P < 0.0002$). (**C**) Line plots showing changes in level of δ1 (left) and δ2 (right) power during wakefulness in light (up) and dark (bottom) phases (SHAM ($n$ = 8) vs Opto-STROKE ($n$ = 9), *two-way ANOVA* with Bonferroni post hoc test. Data are mean ± SEM. $*P < 0.05$, $**P < 0.002$, $***P < 0.0002$). (**D**) Line plots showing changes in level of δ1 (left) and δ2 (right) during NREM in light (up) and dark (bottom) phase (SHAM ($n$ = 8) vs Opto-STROKE ($n$ = 9), *two-way ANOVA* with Bonferroni post hoc test. Data are mean ± SEM. $*P < 0.05$, $**P < 0.002$, $***P < 0.0002$).

*Suppl. Figure 5. Lenzi I. et al*

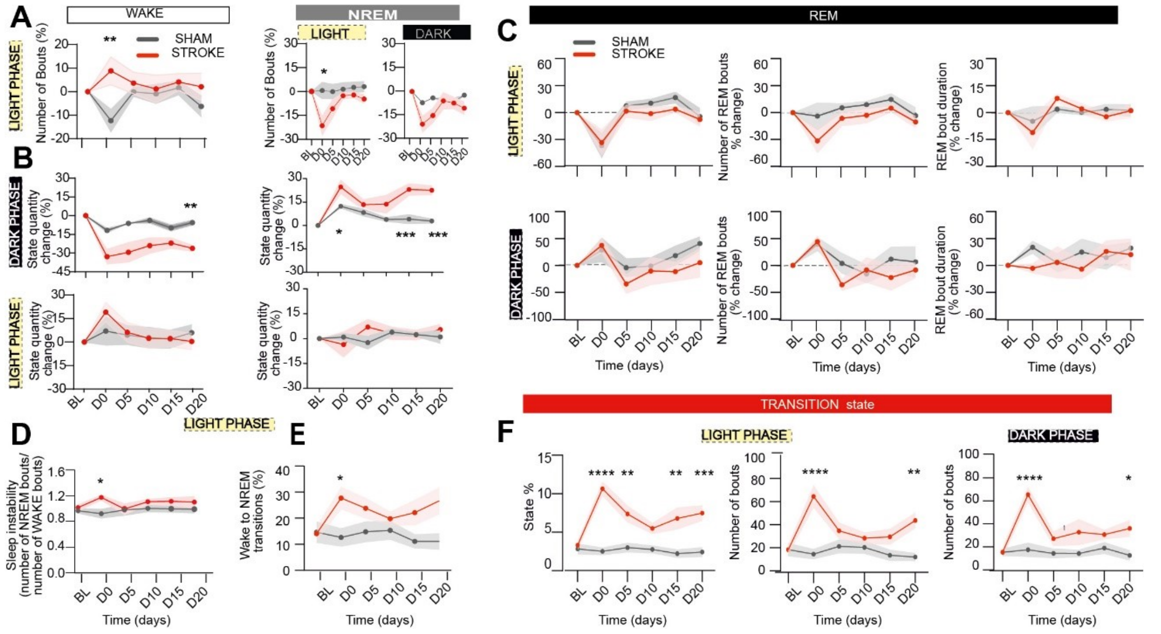


**Supplemental figure 5. Evolution of changes in sleep architecture following IL Opto-STROKE.** (**A**) Line plots showing % change in episodes of wake (left) during the light phase and NREM sleep (right) during the light and dark phases (SHAM (*n* = 10) vs Opto-STROKE (*n* = 9)). (**B**) Line plots showing progression of wake (left) and NREM (right) state quantity % change during the dark (up) and light phase (bottom) (SHAM (*n* = 10) vs Opto-STROKE (*n* = 9)). (**C**) REM state analysis. Left plot: % change over time of: REM state %; central plot: number of REM bouts (middle); right plot: REM mean bout duration (right) (SHAM (*n* = 10) vs Opto-STROKE (*n* = 9)). (**D**) Line plots of over time evolution of sleep instability in the light phase (NREM bouts/WAKE bouts) (SHAM (*n* = 10) vs Opto-STROKE (*n* = 9)). (**E**) Line plots of over time evolution of the number of transitions from wake to NREM sleep during the light phase (SHAM (*n* = 8) vs Opto-STROKE (*n* = 9)). (**F**) Line plots of transitional state analysis: Left plot: over time evolution of transitions % in the light phase; central plot: number of transitions bouts in the light phase; right plot: number of transitions bouts in the dark phase (SHAM (*n* = 10) vs Opto-STROKE (*n* = 9)). Statistical test. *Two-way ANOVA* with Bonferroni post hoc test was used as statistical test, $*P < 0.05$, $**P < 0.002$, $***P < 0.0002$ were consider significant. Data is represented as mean ± SEM.

*Suppl. Figure 6. Lenzi I. et al*

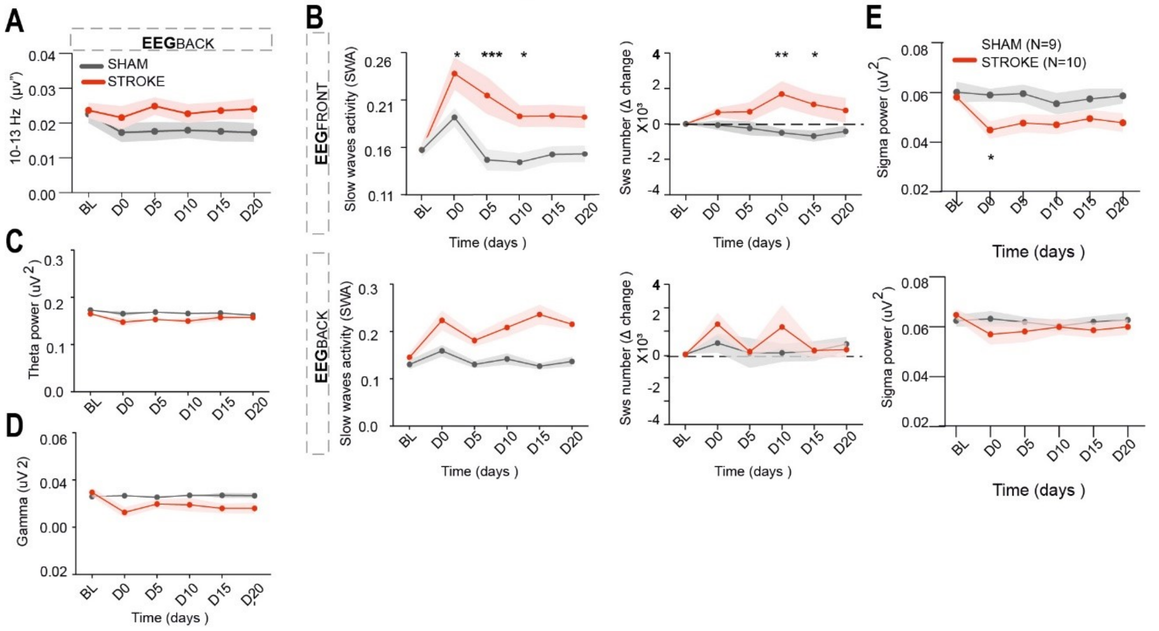


**Supplemental figure 6. Changes in arousability and topography of oscillatory events during wake after CMT Opto-STROKE.** (**A**) From left to right: line plots showing changes over time in slow waves activity (SHAM (*n* = 9) vs Opto-STROKE (*n* = 8)) and number of slow waves in the frontal (up) and parietal (bottom) EEG (SHAM (*n* = 9) vs Opto-STROKE (*n* = 10), *two-way ANOVA with* Bonferroni post hoc test. Data are mean ± SEM. **P* < 0.05, ***P* < 0.002, ****P* <0.0002). (**B**) Line plots showing changes over time in theta power (5-10 Hz) in the frontal (up) and parietal (bottom) EEG (SHAM (*n* = 9) vs Opto-STROKE (*n* = 8), *two-way ANOVA* with Bonferroni post hoc test. Data are mean ± SEM. **P* < 0.05, ***P* < 0.002, ****P* <0.0002). (**C**) Line plots showing changes over time in alpha power (10-13 Hz) in the parietal EEG (SHAM (*n* = 9) vs Opto-STROKE (*n* = 8), *two-way ANOVA* with Bonferroni post hoc test. Data are mean ± SEM. *p < 0.05). (**D)** Line plots showing changes over time in gamma power in the parietal EEG (SHAM (*n* = 8) vs Opto-STROKE (*n* = 10), *two-way ANOVA* with Bonferroni post hoc test. Data are mean ± SEM. **P* < 0.05, ***P* < 0.002, ****P* <0.0002).

*Suppl. Figure 7 Lenzi I. et al*

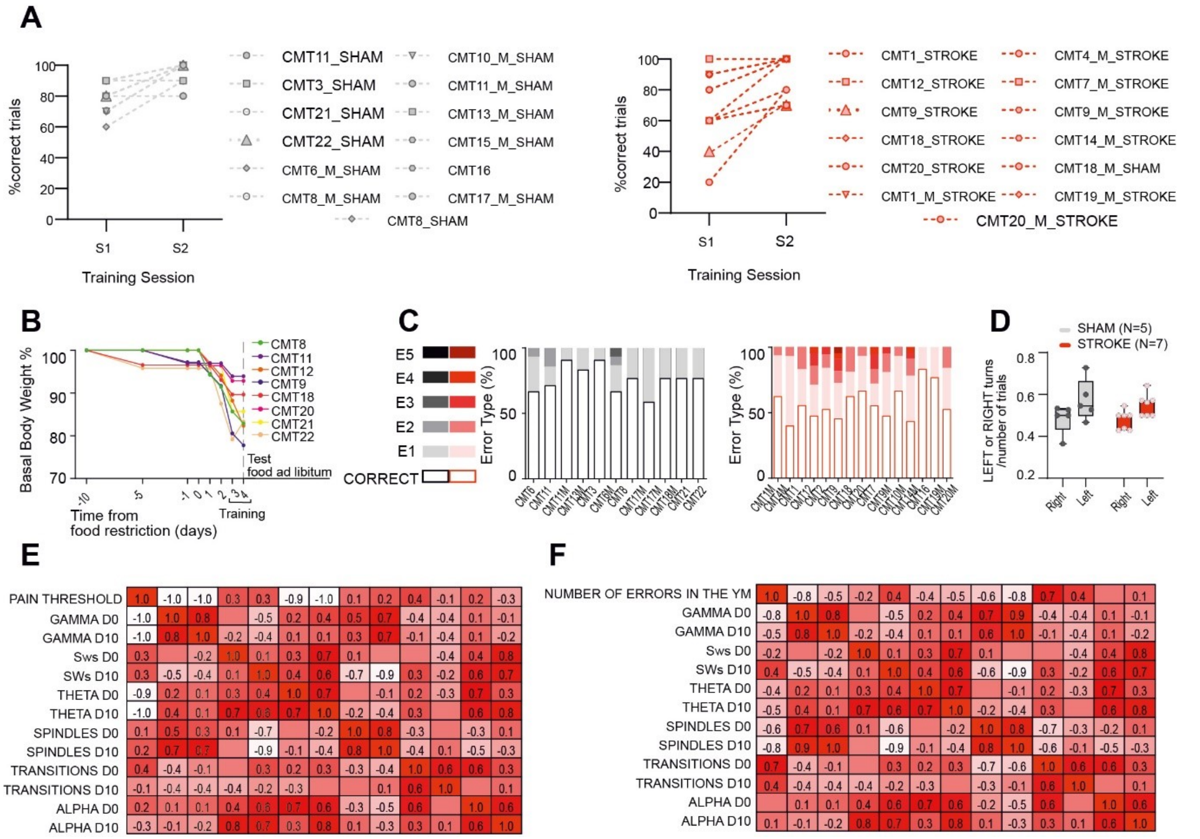


**Supplemental figure 7. Individual data from IL Opto-STROKE and SHAM animals in the Y-maze alternating working memory task.** (**A**) Curves showing body weight (BW) monitoring during food restriction protocol (exclusion-threshold at 80 % of basal BW). (**B**) Individual performance of SHAM (grey, left) and Opto-STROKE (red, right) animals in the YM session (Error 1: E1; error 2: E1; error 3: E3; error 4: E4; error 5: E5) as a fraction of cumulative performance (Total trials = 100%), with left legend indicating type of errors. (**C**) Minimum-maximum box-plots showing number of left and right turns performed by SHAM and Opto-STROKE (SHAM ($n$ = 5) vs Opto-STROKE ($n$ = 7), unpaired *t-test*, **P* < 0.05, ***P* < 0.002, ****P* <0.0002). (**D**) Learning curves over the two training sessions (S1 and S2) of SHAM (left) and Opto-STROKE animals (right) (Data are presented as raw values of individual animal % correct responses over a training session). (**D**) Correlation analysis between pain sensitivity threshold with sleep transitions and sleep oscillations at day 0 and 10 after Opto-stroke induction. (**F**) Correlation analysis between errors performed at the day of testing in the Y-maze working memory task with sleep transitions and sleep oscillations at day 0 and 10 after Opto-STROKE induction (*Pearson Correlation*. Data are: -1 < r < +1 (white= min value; red= max value).

### Supplemental Table 1.

| Transitional States % | Light phase | | |
|---|---|---|---|
| | Mean SHAM ± SD | Mean Opto-STROKE ± SD | Two-way ANOVA *P* value SHAM vs Opto-STROKE |
| **D0** | 2.501 ± 1.503 | 10.648 ± 3.185 | $P < 0.001$ |
| **D5** | 2.995 ± 1.885 | 7.367 ± 3.612 | $P = 0.002$ |
| **D10** | 2.750 ± 1.463 | 5.478 ± 2.129 | $P = 0.19$ |
| **D15** | 2.208 ± 1.567 | 6.779 ± 4.281 | $P = 0.001$ |
| **D20** | 2.423 ± 1.857 | 7.478 ± 3.046 | $P = 0.01$ |

### Supplemental Table 2.

| EEG topography | State | SWA | | | Delta1 | | | Delta2 | | |
|---|---|---|---|---|---|---|---|---|---|---|
| | | Mean Sham ± SD | Mean Stroke ± SD | Two-way ANOVA p-value sham vs stroke | Mean Sham ± SD | Mean Stroke ± SD | Two-way ANOVA p-value sham vs stroke | Mean Sham ± SD | Mean Stroke ± SD | Two-way ANOVA p-value sham vs stroke |
| $EEG^{FRONT}$ | ALL | -8.45± 4.51 | 18.36± 11.43 | F (1, 82) = 16.09 P=0.0001 | -2.03± 4.25 | 9.44± 11.47 | F (1, 90) = 3.288 P=0.0731 | 0.11± 0.08 | 0.46± 0.26 | F (1, 93) = 12.27 P=0.0007 |
| | WAKE | 0.16± 0.01 | 0.20± 0.02 | F (1, 78) = 25.83 P<0.0001 | 0.17± 0.10 | 0.09± 0.11 | F (1, 102) = 0.5849 P=0.4462 | 0.02± 0.03 | 0.48± 0.31 | F (1, 78) = 25.43 P<0.0001 |
| | NREM | 0.24± 0.00 | 0.26± 0.01 | F (1, 67) = 8.062 P=0.0060 | -0.02± 0.04 | -0.03± 0.10 | F (1, 81) = 0.9910 P=0.3225 | -0.13± 0.08 | 0.02± 0.05 | F (1, 88) = 9.933 P=0.0022 |
| $EEG^{PAR}$ | ALL | -0.61± 6.95 | 0.83± 6.78 | F (1, 97) = 0.04716 P=0.8285 | 5.56± 7.36 | 5.04± 9.81 | F (1, 90) = 0.004922 P=0.9442 | -0.24± 7.22 | -9.63± 9.01 | F (1, 86) = 2.247 P=0. 1375 |
| | WAKE | 0.14± 0.01 | 0.16± 0.02 | F (1, 86) = 3.261 P=0.0745 | 5.85± 7.67 | 0.42± 7.64 | F (1, 96) = 0.4711 P=0.4941 | -8.37± 8.39 | 4.11± 5.66 | F (1, 90) = 3.444 P=0.0668 |
| | NREM | 0.21± 0.01 | 0.21± 0.01 | F (1, 63) = 9.399e-005 P=0.9923 | 2.17± 6.02 | -2.95± 7.35 | F (1, 89) = 0.7138 P=0.4004 | -6.13± 4.80 | 1.01± 3.93 | F (1, 79) = 5.338 P=0.0235 |

### Supplemental Table 3.

| EEG topography | State | SWA | | | Delta1 | | | Delta2 | | |
|---|---|---|---|---|---|---|---|---|---|---|
| | | Mean Sham ± SD | Mean Stroke ± SD | Two-way ANOVA p-value sham vs stroke | Mean Sham ± SD | Mean Stroke ± SD | Two-way ANOVA p-value sham vs stroke | Mean Sham ± SD | Mean Stroke ± SD | Two-way ANOVA p-value sham vs stroke |
| $EEG^{FRONT}$ | ALL | -6.00± 3.58 | 8.37± 5.27 | F (1, 84) = 15.41 P=0.0002 | -2.05± 3.29 | -1.63± 9.23 | F (1, 86) = 0.008148 P=0.9283 | -16.54± 8.27 | 18.96± 16.45 | F (1, 83) = 17.73 P<0.0001 |
| | WAKE | 0.16± 0.017 | 0.20± 0.027 | F (1, 73) = 38.75 P<0.0001 | -0.05± 0.07 | 0.03± 0.11 | F (1, 92) = 1.204 P=0.2753 | -0.18± 0.09 | 0.21± 0.18 | F (1, 91) = 12.39 P=0.0007 |
| | NREM | -0.15± 0.08 | 0.22± 0.16 | F (1, 73) = 2.912 P=0.0922 | -0.10± 0.06 | 0.01± 0.07 | F (1, 75) = 3.617 P=0.0610 | -0.15± 0.08 | 0.22± 0.16 | F (1, 75) = 14.39 P=0.0003 |
| $EEG^{PAR}$ | ALL | -3.75± 2.56 | -2.34± 8.24 | F (1, 102) = 0.06462 P=0.7998 | 3.20± 5.49 | -0.71± 8.10 | F (1, 103) = 0.3015 P=0.5841 | -12.96± 7.07 | -4.58± 5.70 | F (1, 103) = 2.433 P=0.1219 |
| | WAKE | 0.14± 0.01 | 0.19± 0.03 | F (1, 85) = 26.88 P<0.0001 | 9.93± 9.93 | -5.34± 6.68 | F (1, 95) = 3.242 P=0.0750 | -9.90± 5.27 | 1.72± 6.71 | F (1, 85) = 3.548 P=0.0630 |
| | NREM | 0.22± 0.00 | 0.22± 0.01 | F (1, 82) = 0.3566 P=0.5521 | 1.12± 5.76 | -2.90± 5.58 | F (1, 99) = 0.4105 P=0.5232 | -0.08± 5.68 | -10.55± 5.24 | F (1, 93) = 4.206 P=0.0431 |

### Supplemental Table 4.

| Response type % in the YM | Mean SHAM ± SD | Mean Opto-STROKE ± SD | Two-way ANOVA *P*-values SHAM vs Opto-STROKE |
|---|---|---|---|
| **Correct Response** | 76.03 ± 3.00 | 57.28 ±3.20 | $P < 0.0001$ |
| **Type1 Error** | 20.85 ±2.70 | 28.22 ± 2.33 | $P = 0.03$ |
| **Type2 Error** | 2.51 ±1.42 | 9.99 ±1.46 | $P = 0.03$ |
